# Convergent and Divergent Brain–Cognition Development

**DOI:** 10.1101/2025.06.06.658294

**Authors:** Yapei Xie, Shaoshi Zhang, Csaba Orban, Leon Qi Rong Ooi, Ru Kong, Dorothea L. Floris, Xi-Nian Zuo, Elvisha Dhamala, Avram J Holmes, Lucina Q. Uddin, Thomas E Nichols, Adriana Di Martino, B.T. Thomas Yeo

## Abstract

How brain networks and cognition co-evolve during development remains poorly understood. Here, we use resting-state functional magnetic resonance imaging (rs-fMRI) and cognitive data at baseline and Year 2 of 2,949 individuals in the Adolescent Brain Cognitive Development (ABCD) Study to examine how stable and changing features of brain network organization predict cognitive development during early adolescence. We find that baseline resting-state functional connectivity (FC) more strongly predicts future cognitive ability than baseline cognitive ability. Models trained on baseline FC to predict baseline cognition generalize better to Year 2 FC and cognition, suggesting that brain–cognition relationships strengthen over time.

Intriguingly, baseline FC outperforms longitudinal FC change in predicting future cognitive ability. One potential reason is the lower reliability of FC change compared to baseline FC: ICC = 0.24 vs. 0.56. However, reducing baseline FC’s reliability by shortening scan duration only partially narrows the predictive gap, suggesting reliability alone cannot be the full explanation. Furthermore, neither baseline FC nor FC change meaningfully predicts longitudinal change in cognitive ability. We also identify converging and diverging predictive network features across cross-sectional and longitudinal models of brain–cognition relationships, revealing a multivariate twist on Simpson’s paradox. Together, these findings suggest that during early adolescence, stable individual differences in brain functional network organization play a more critical role than dynamic changes in shaping future cognitive outcomes.

## INTRODUCTION

A major goal in cognitive neuroscience is to understand how individual differences in brain network development relate to variability in cognitive development. This is particularly salient during the transition from childhood to adolescence, a period marked by a shift from concrete to more abstract and logical thinking, which supports the growing ability to manage complex tasks and navigate increasingly demanding environments (Piaget, 1952; Luna et al., 2004; Shaffer, 2013; Gopnik et al., 2017). Here, we leverage longitudinal resting-state fMRI and cognitive data from 2,949 children (ages 8.9 to 13.5) to examine how individual-level brain–cognition predictive models develop longitudinally during this critical period.

Resting-state fMRI (rs-fMRI) is a powerful tool for examining functional brain network organization (Biswal et al., 1995; Fox et al., 2006; Smith et al., 2009; Yeo et al., 2011). Previous studies have developed cross-sectional brain–cognition models that can predict individual-level cognitive performance in children, adolescents, and young adults based on resting-state functional connectivity (Finn et al., 2015; Rosenberg et al., 2016; Kong et al., 2019; Cui et al., 2020; Sripada et al., 2020; Chen et al., 2022; Keller et al., 2023; Zhi et al., 2024). While valuable, these cross-sectional models reflect a single snapshot of brain–cognition relationships, neglecting dynamic changes that can only be revealed in longitudinal designs (Guillaume et al., 2014; Sorensen, Walhovd, & Fjell, 2021; Kang et al., 2024; Saha et al., 2024).

Conceptually, cross-sectional analyses reveal between-individual traits, which might differ from dynamic within-individual changes captured by longitudinal designs (Curran & Bauer, 2011; Walhovd, Lovden, & Fjell, 2023). Figure 1a illustrates the classical example of how cross-sectional (between-individual) and longitudinal (within-individual) estimates can diverge for a single phenotype, a phenomenon known as the Simpson’s paradox (Pearl, 2014; Walhovd et al., 2017; Di Biase et al., 2023). When examining the interplay between two phenotypes, such as brain and cognition, these divergences can become even more complex.

**Figure 1.**
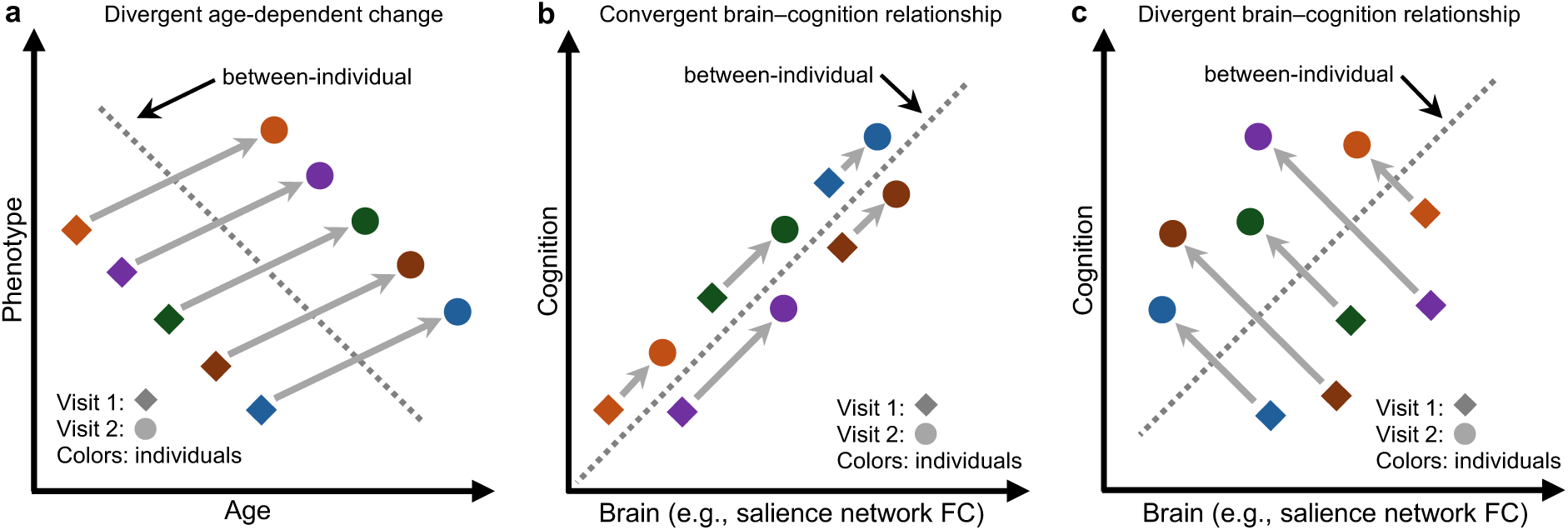
Illustration of how cross-sectional and longitudinal estimates can converge or diverge. (a) Classical (Simpson’s paradox) illustration, where within-individual changes (arrows) diverge from between-individual differences (dashed line). (b) Convergent cross-sectional and longitudinal brain-cognition models. For example, greater salience network connectivity predicts better cognitive ability among children cross-sectionally (dashed line). Assuming a causal relationship, individuals with larger increases in salience network FC should enjoy greater cognitive gains longitudinally (arrows). (c) Divergent cross-sectional and longitudinal brain-cognition models. Similar to panel (b), greater salience network connectivity predicts better cognitive ability among children cross-sectionally (dashed line). However, individuals with larger reductions in salience network FC enjoy greater cognitive gains longitudinally (arrows). Figure S1 illustrates other possible divergences between cross-sectional and longitudinal estimates of brain–cognition relationship. We hypothesize that convergence and divergence vary across brain networks.

To illustrate the distinction between cross-sectional and longitudinal brain–cognition models, consider a simplified scenario. Imagine Luca is a cognitively advanced child whose brain connectivity patterns differ from his peers. As his peers grow older and become more cognitively capable, do their brains become more like Luca’s? If so, this suggests convergence between cross-sectional and longitudinal models. If not – if children who make the greatest cognitive gains develop different connectivity patterns from Luca – this indicates divergence.

These contrasting scenarios are critical for understanding individual-level prediction. For example, a previous cross-sectional ABCD study found that stronger salience network connectivity predicted better cognitive ability at the individual level (Chen et al., 2022). If this relationship reflects a causal mechanism, one would expect that children with greater increases in salience network connectivity over time would also show greater cognitive improvement (Figure 1b). However, cross-sectional and longitudinal models may not always align: it is also possible that children with larger reductions in salience connectivity exhibit the most cognitive gains (Figure 1c). While we use the salience network here as an example, we hypothesize that patterns of convergence and divergence in predictive features will vary across brain networks.

Using longitudinal rs-fMRI and cognitive data collected at baseline and Year 2 from 2949 participants of the ABCD Study (Casey et al., 2018; Jernigan, Brown, & Dowling, 2018; Volkow et al., 2018), we investigated how brain functional connectivity (FC) and cognition co-evolve during early adolescence. Although group-level cognitive performance improves over time, individual differences in cognition remain highly stable. Similarly, FC shows both longitudinal change and individual stability, organized along the sensory–association (S-A) axis: sensory regions change more over time but are less stable across individuals, while association regions show the opposite pattern. Focusing on brain–cognition development, we find that baseline FC is a stronger predictor of future cognitive ability than baseline cognition. Moreover, models trained on baseline FC to predict baseline cognition perform better when applied to data at Year 2. These results suggest that brain–cognition relationships strengthen over time. Intriguingly, longitudinal FC change is a weaker predictor of future cognition than baseline FC, even after accounting for differences in reliability. Finally, neither baseline FC nor FC change meaningfully predicts changes in cognition over two years. We also identify convergent (Figure 1b) and divergent (Figure 1c) network features across cross-sectional and longitudinal models. Together, these findings suggest that while brain–cognition relationships are refined over development, they are shaped most strongly by stable individual differences in brain network organization.

## RESULTS

### Overview

We first investigated individual differences in cognitive change and FC change across the two ABCD Study timepoints. We then explored cross-sectional FC–cognition relationship. Finally, we examined the longitudinal relationship between FC and cognition.

### Individual differences in longitudinal cognitive and FC changes

We considered a sample of 2949 participants from the ABCD Study (releases 4 and 5), who had seven cognitive measures and rs-fMRI data that survived quality control at both baseline and Year 2 (Table 1). We also computed the first principal component (PC1) of the seven cognitive measures.

**Table 1.**
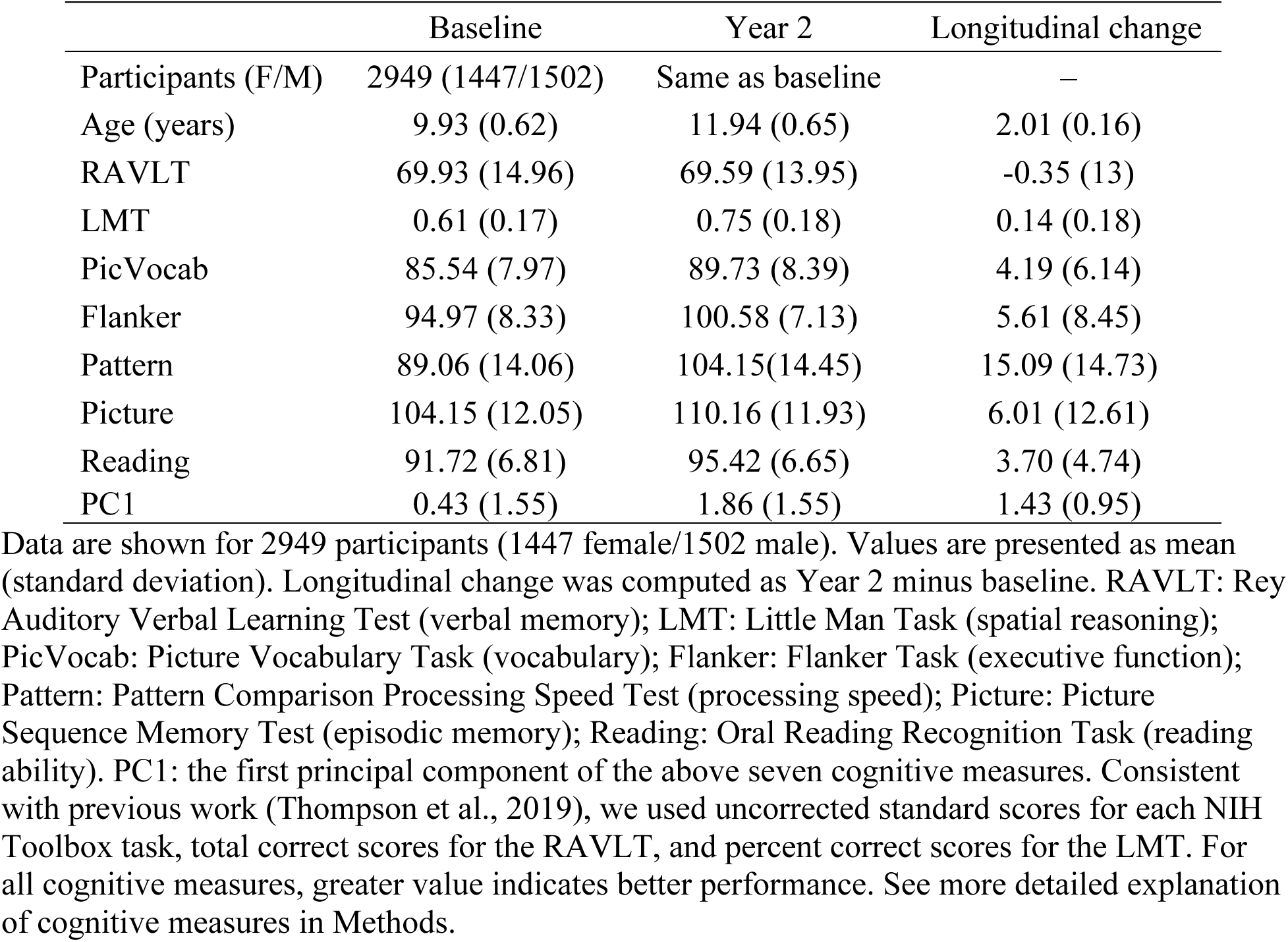
Age and cognitive measures at baseline, Year 2, and their longitudinal change.

Cognitive ability at baseline and Year 2 was positively correlated, suggesting that individuals with higher baseline cognition generally maintained their cognitive advantage over their peers at Year 2 (Figure 2a). Although participants generally showed improved cognitive ability from baseline to Year 2, except for RAVLT (Figure 2b), which was consistent with a previous study (Anokhin et al., 2022), substantial individual variability in longitudinal cognitive change was observed across most cognitive measures (Figure 2c).

**Figure 2.**
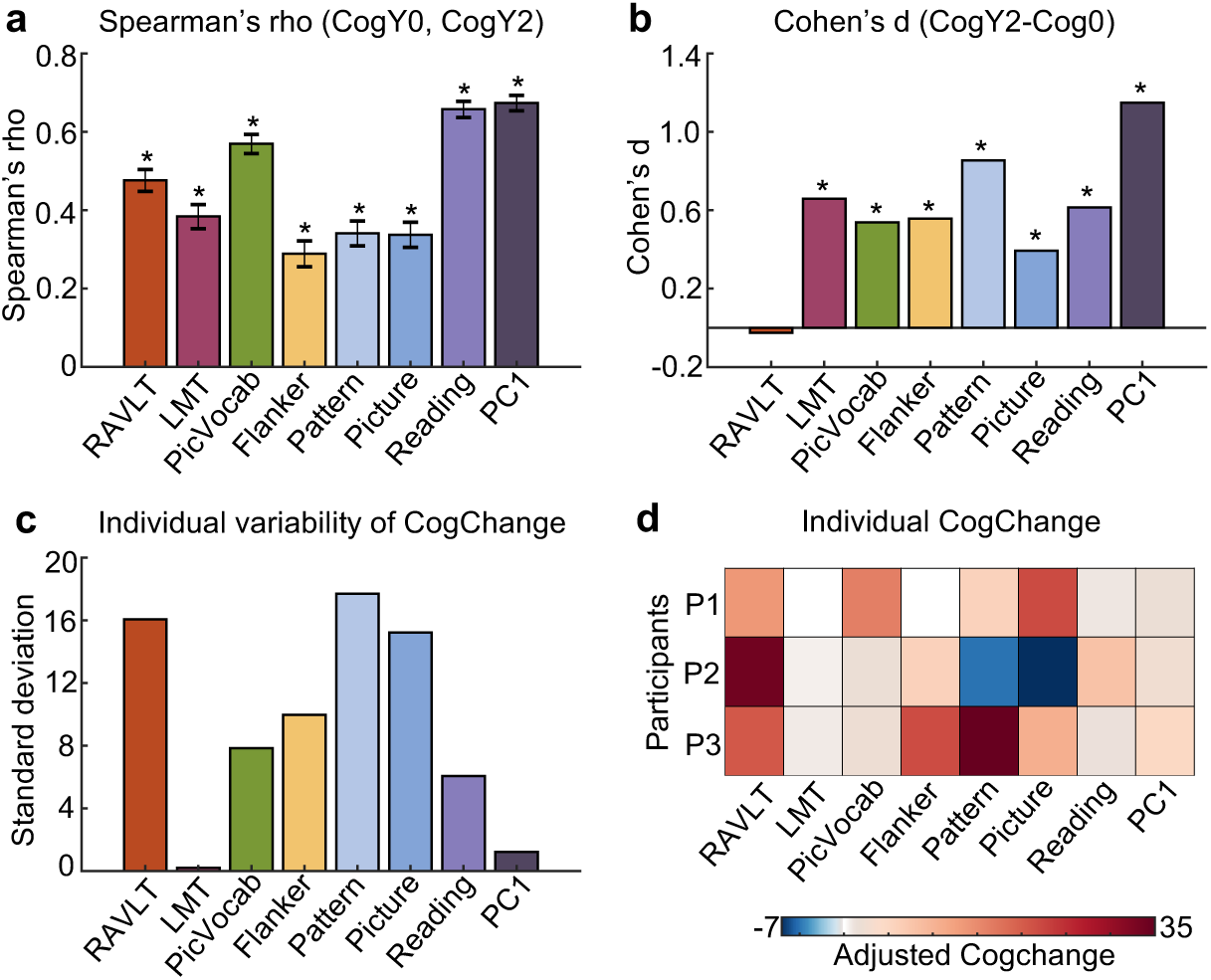
Individual differences in longitudinal cognitive change during the transition from childhood to adolescence. (a) Spearman’s correlation between baseline and Year 2 cognitive measures. The positive correlations indicate that children with higher baseline cognition generally maintained their cognitive advantage over their peers at Year 2. (b) Longitudinal cognitive change at the group level estimated from a linear mixed effects model. (c) Individual variability in longitudinal cognition change. For each individual and each cognitive measure, cognition change was defined as the difference in scores between the two timepoints. Sex, baseline age and age interval (between baseline and Year 2) were regressed out. Standard deviation was then computed across individuals. (d) Longitudinal changes of the eight cognitive measures for three participants. RAVLT: Rey Auditory Verbal Learning Test (verbal memory); LMT: Little Man Task (spatial reasoning); PicVocab: Picture Vocabulary Task (vocabulary); Flanker: Flanker Task (executive function); Pattern: Pattern Comparison Processing Speed Test (processing speed); Picture: Picture Sequence Memory Test (episodic memory); Reading: Oral Reading Recognition Task (reading ability). PC1: the first principal component of the above seven cognitive measures. Asterisks (*) indicate statistical significance after false discovery rate (FDR) correction at q < 0.05 (Benjamini & Hochberg, 1995).

As an example, Figure 2d shows the longitudinal changes of the eight cognitive measures for three participants. The Pearson’s correlations among the three cognitive change profiles were small (average r = 1.9 × 10⁻^4^), suggesting that the patterns of cognitive improvement were highly different across the three participants. For example, compared with their peers, participant 2 became substantially worse in the Picture Sequence Memory Task (“Picture”), but exhibited relatively strong improvement in the Rey Auditory Verbal Learning Task (RAVLT). On the other hand, participant 1 showed strong improvements in both Picture and RAVLT tasks.

Positive correlations indicate that children exhibiting stronger brain connectivity at baseline continued to exhibit stronger brain connectivity than their peers at Year 2. 99.995% of entries were significant after FDR correction with q < 0.05. (d) Visualization of FC stability at the regional level, by summing the rows of panel (c). (e) Longitudinal FC change at the group level based on a linear mixed effects model. 54.96% of entries were significant after FDR correction with q < 0.05. (f) Visualization of longitudinal FC change at the regional level, by summing the rows of panel (e). (g) Individual variability in longitudinal FC change. FC change (z value) was computed for each FC edge (Afyouni, Smith, & Nichols, 2019). Sex, baseline age, age interval (between baseline and Year 2) and head motion at two timepoints were regressed out. Standard deviation was then computed across individuals. (h) Visualization of individual variability in longitudinal FC change at the regional level, by summing the rows of panel (g). (i-k) Individual level FC change for three participants.

The rs-fMRI was used to compute a 419 × 419 FC matrix for each participant and each time point on the 400-region Yan parcellation (Yan et al., 2023) (Figure 3a) and 19 subcortical regions (Fischl et al., 2002) (Figure 3b). Similar to cognition, baseline and Year 2 FC was positively correlated, suggesting that children exhibiting stronger brain connectivity at baseline continued to exhibit stronger brain connectivity than their peers at Year 2 (Figure 3c). By averaging the rows of Figure 3c, clear regional variation was observed along the sensory-association (S-A) axis, with heteromodal association cortex being the most stable, while sensory-motor and visual systems were the least stable (Figure 3d). An inverse pattern emerged in longitudinal FC change: sensory-motor and visual systems showed the greatest longitudinal change, while heteromodal association cortex exhibited the least change (Figures 3e and 3f).

**Figure 3.**
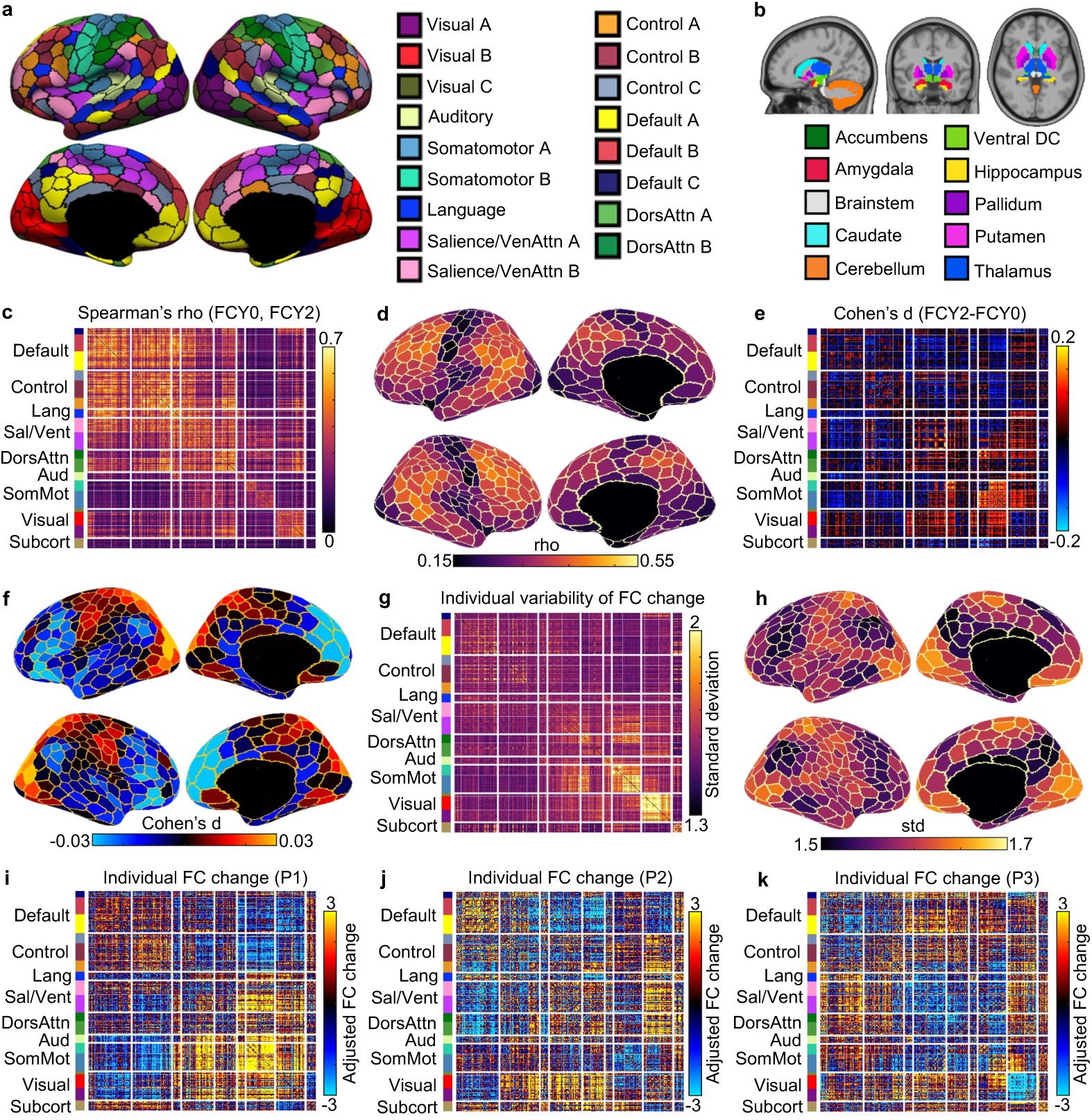
Individual differences in longitudinal functional connectivity (FC) change. (a) Cortical parcellation of 400 regions (Yan et al., 2023), which is the homotopic variant of the Schaefer parcellation (Schaefer et al., 2018). Parcel colors are assigned corresponding to 17 large-scale networks (Kong et al., 2021). (b) 19 subcortical regions (Fischl et al., 2002). 419 × 419 FC matrices were computed based on the 419 cortical and subcortical regions. (c) Spearman’s correlation (stability) between baseline (FCY0) and Year 2 FC (FCY2) for each FC edge.

However, we also observed notable individual differences in FC change (Figure 3g), especially for somatomotor and visual networks, as well as medial prefrontal cortex (Figure 3h). As an example, Figures 3i to 3k show the longitudinal FC changes of three participants. The Pearson’s correlations among the three FC change profiles were small (average r =-0.04), suggesting that the patterns of FC change were highly different across the three participants. For example, relative to their peers, participant 1 exhibited strong increases in FC within the somatomotor network, in contrast to participant 2 who exhibited strong decreases in somatomotor FC.

The substantial individual differences in cognition and FC change motivate the study of how the brain–cognition relationship evolves during transition from childhood to adolescence.

### The relationship between FC and cognition strengthens with development

We next examined cross-sectional relationships between FC and cognition. We first used kernel ridge regression (KRR) to predict cross-sectional cognition from cross-sectional FC. This was achieved via a nested cross-validation procedure that was repeated 120 times for robustness (see methods for details). Care was taken so that participants from the same site were not split between training and test sets, so prediction performance was always measured in out-of-sample sites not used to train the models.

Consistent with previous work (Chen et al., 2022; Zhi et al., 2024), baseline FC was able to predict all eight cognitive measures at baseline (FDR < 0.05), with prediction accuracy of r = 0.43 for the first cognitive principal component PC1 (Figure 4a). Baseline FC could also effectively predict all eight cognitive measures at Year 2, with accuracy of r = 0.46 for PC1. We additionally found that Year 2 FC predicted all cognitive measures at Year 2, achieving accuracy of r = 0.49 for PC1.

**Figure 4.**
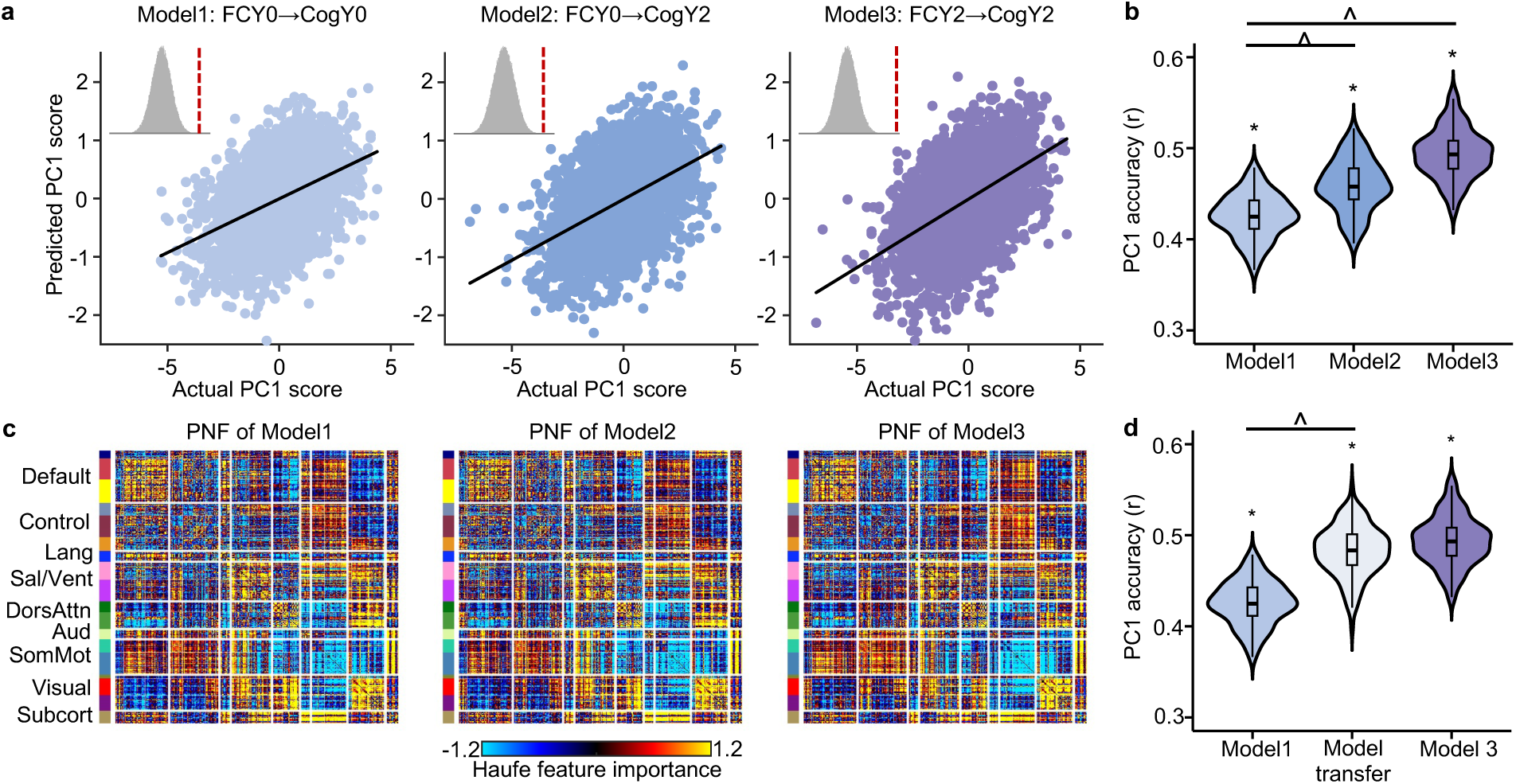
Enhanced FC–cognition relationships during development. (a) Correlation between the actual cognitive principal component (PC1) score and the predicted PC1 score. Model 1 predicted baseline cognition using baseline FC (FCY0 → CogY0). Model 2 predicted Year 2 cognition using baseline FC (FCY0 → CogY2). Model 3 predicted Year 2 cognition using Year 2 FC (FCY2 → CogY2). Insets show null distributions based on 1000 permutations with red dash line corresponding to actual prediction accuracy. (b) Comparison of prediction accuracies across the three models. Each value in the violin plot represents the accuracy (r) for a single cross-validation fold. Asterisks (*) denote above chance prediction after multiple comparisons correction (FDR q < 0.05). Carets (^) denote statistically significant differences between models based on the corrected resampled t-test (Nadeau & Bengio, 2003) with FDR q < 0.05. (c) Predictive network feature (PNF) matrix for each model. PNF was computed using the Haufe transformation (Haufe et al., 2014). Positive values indicate that higher FC were associated with higher predicted cognitive scores, while negative values indicate higher FC were associated with lower predicted cognitive scores. For visualization purposes, each predictive network feature matrix was normalized by dividing all values by the standard deviation of the entire matrix. (d) Models trained on baseline FC to predict baseline cognition improve in accuracy when applied to Year 2 FC and Year 2 cognition. Model1 was used to predict Y2 cognition from Year 2 FC, which we refer to as “model transfer”. Figures S2 and S3 repeat panels (b) and (d) for the other seven cognitive measures, respectively. FC: functional connectivity; PC1: the first principal component of the seven cognitive measures.

Year 2 FC’s prediction of Year 2 cognition was statistically better than baseline FC’s prediction of baseline cognition, and baseline FC predicted Year 2 cognition more accurately than it predicted cognition at baseline (FDR < 0.05; Figure 4b). Figure S2 showed the comparisons for the other seven cognitive measures. These findings collectively suggest enhanced FC–cognition relationships during development.

We further utilized Haufe transformation (Haufe et al., 2014) to interpret each predictive model. Briefly, the covariance between each FC edge and the predicted cognition score was computed across participants. This analysis produced a 419 × 419 predictive network feature (PNF) matrix for each model and cognitive measure, where positive values indicate that higher FC were associated with higher predicted cognition scores. Previous studies have demonstrated that the Haufe transformation led to more valid interpretation of predictive models (Haufe et al., 2014), which are highly reliable and generalizable across different predictive algorithms (Tian & Zalesky, 2021; Chen et al., 2022; Chen et al., 2023).

Predictive network features were highly similar across all three models, with positive and negative contributions widely distributed across the entire connectome (Figure 4c). The predictive network features of Model 3 (i.e., FCY2→CogY2) exhibited a more pronounced contrast (Figure 4c), with both stronger positive and stronger negative contributions compared to predictive network features of Model 1 (i.e., FCY0→CogY0) and Model 2 (i.e., FCY0→CogY2).

### Models trained on baseline FC to predict baseline cognition improve in accuracy when applied to Year 2 FC and Year 2 cognition

Given the high similarities across the PNF matrices of all three models (Figure 4c), we hypothesized that the model trained on baseline data would perform well when applied to Year 2 data. Surprisingly, we found that the model trained on baseline FC to predict baseline cognition showed improved accuracy when applied to predict Year 2 cognition from Year 2 FC for PC1 (FDR q < 0.05; Figure 4d). This result suggests that the multivariate relationship between brain and cognition is similar at baseline and Year 2, but the strength of the relationship increases over time, yielding better prediction accuracy. Therefore, early established functional architecture provides a foundation that becomes increasingly effective in supporting cognitive ability. Figure S3 showed the comparisons for the other seven cognitive measures. To ensure that our results are not influenced by lower head motion in Year 2 compared to baseline, we repeated the analysis in this section using a subset of participants with comparable motion levels across the two time points (p = 0.18), which yielded similar conclusions (Figure S4).

### Baseline FC is more predictive of cognition at Year 2 than longitudinal FC changes even accounting for reliability differences

So far, we have demonstrated that there is a strong cross-sectional relationship between FC and cognition, and the relationship strengthens with development (Figure 4). We hypothesized that longitudinal FC change might also predict future cognitive outcomes.

Longitudinal FC change was able to predict all cognitive measures (except for the Pattern Comparison Processing Speed Task) at Year 2 (FDR q < 0.05; Figures 5a and 5b, Figure S5). However, the best prediction accuracy was only 0.13 (for PC1). In general, baseline FC outperformed longitudinal FC change in predicting all cognitive measures at Year 2 (FDR q < 0.05; Figures 5a, 5b and S5). Predictive network features (PNFs) for PC1 were modestly similar across the two models (r = 0.33, pspin < 0.001). Similar conclusions were obtained (Figure S6), when using rate of FC change between Year 2 and baseline (instead of FC change) to predict cognitive measures at Year 2.

**Figure 5.**
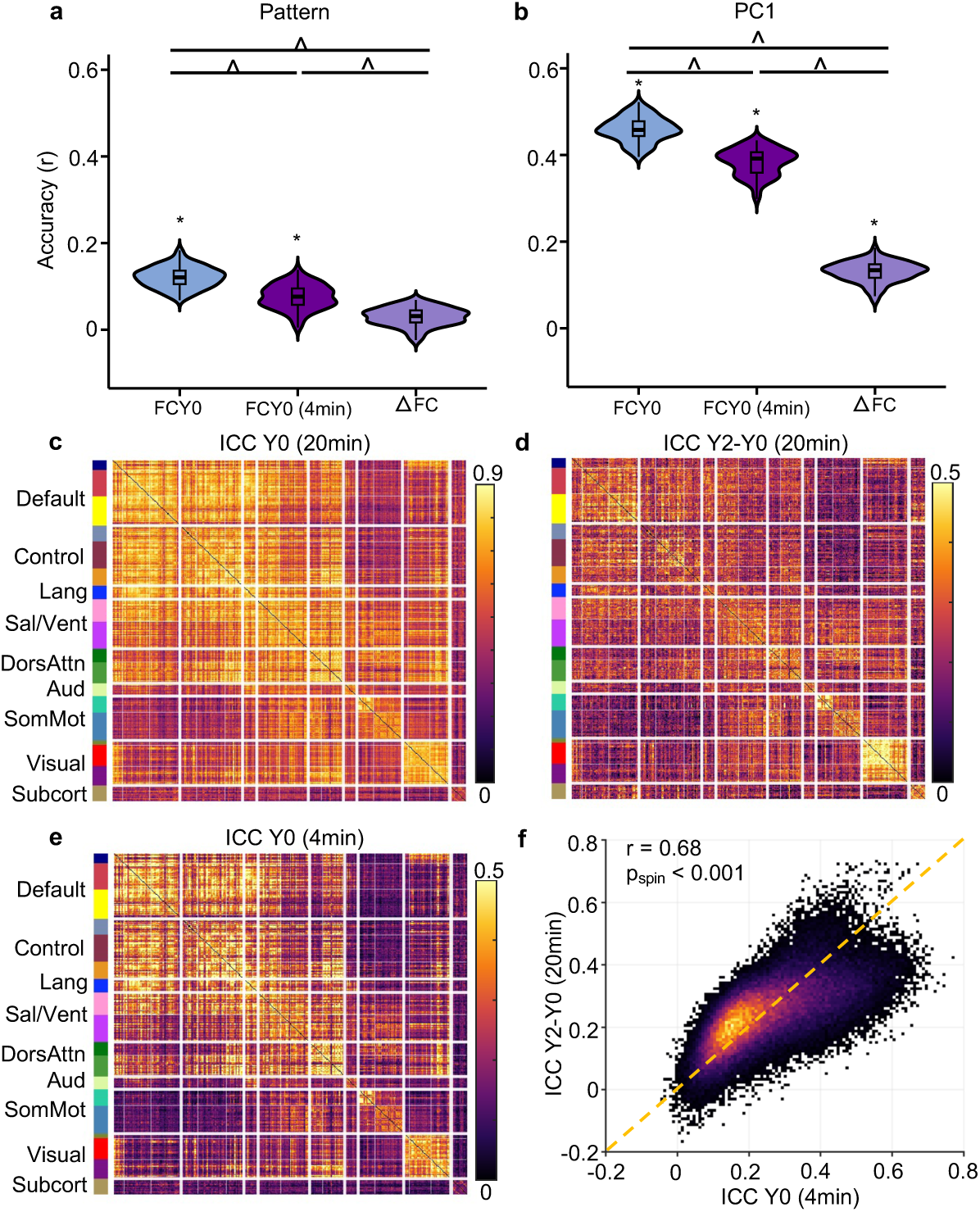
Cross-sectional baseline FC outperforms longitudinal FC change in predicting cognition at Year 2 even accounting for reliability differences. (a) Comparison of prediction accuracies for Pattern Comparison Processing Speed Test at Year 2 across three models. “FCY0” refers to the model trained on baseline FC to predict Year 2 cognition (FCY0 → CogY2). “FCY0 (4min)” uses baseline FC computed from the first 4 minutes of fMRI data to predict Year 2 cognition (FCY0 (4min) → CogY2). “ΔFC” employs the FC change (between Year 2 and baseline) to predict cognition at Year 2 (ΔFC→ CogY2). (b) Same as panel (a) but for PC1. Figure S5 repeats panels (a) and (b) for other 6 cognitive measures. Asterisks (*) denote above chance prediction after multiple comparisons correction (FDR q < 0.05). Carets (^) denote statistically significant difference between models based on the corrected resampled t-test (FDR q < 0.05). (c) Estimated intra-class correlation (ICC) for baseline FC based on 20 minutes of fMRI data. (d) Estimated ICC for FC change based on 20 minutes of fMRI data. (e) Estimated ICC for baseline FC based on 4 minutes of fMRI data. (f) Comparable ICCs were observed for 4-minute baseline FC and 20-minute FC change. Each dot represents the ICC for an FC edge, computed from baseline (x-axis) and FC change (y-axis); dot color indicates point density, with warmer colors reflecting a higher concentration of edges. ICCs were strongly positively correlated (r = 0.68 without subcortical regions; r = 0.67 with subcortical regions). Statistical significance was determined using a spatial permutation (“spin”) test (see Methods) (Alexander-Bloch et al., 2018; Vasa et al., 2018). The yellow dashed line denotes the identity line (y = x) for reference.

The lower predictive accuracy for longitudinal FC change might be due to the lower reliability of FC change compared with cross-sectional FC. We can mathematically express the reliability of FC change in terms of the reliability of baseline FC, reliability of Year 2 FC, and the similarity between the FC of the two timepoints (Equation 4 in Methods). Indeed, the high FC stability between baseline and Year 2 (Figure 3c) is detrimental to the reliability of FC change.

To estimate the reliabilities of baseline FC (Figure 5c) and longitudinal FC change (Figure 5d), we split the four resting-state fMRI runs from each participant into two groups of two runs each, treating them as test and retest sessions. A formula relating FC reliability and scan duration (Ooi et al., 2024) was then used to extrapolate the reliability estimates to 20 minutes. Since both test and retest data were acquired within the same session, these estimates likely represent an upper bound on true test-retest reliability.

As expected, the reliability of baseline FC (Figure 5c) was substantially higher than the reliability of longitudinal FC change (Figure 5d): mean of 0.56 vs 0.24, respectively. To assess whether lower reliability might account for the weaker prediction performance of longitudinal FC change, we artificially shortened the fMRI scan duration used to compute baseline FC from 20 minutes to 4 minutes. The resulting reliabilities were now similar between the 4-minute baseline FC and the longitudinal FC change (Figures 5d, 5e and 5f), with both showing a mean of 0.24 and a correlation of 0.68 (pspin < 0.001).

Unsurprisingly, 4-minute baseline FC generally showed lower predictive performance than 20-minute baseline FC, but was still better than longitudinal FC change (FDR < 0.05; Figures 5a, 5b and S5). This implies that the lower prediction accuracy of longitudinal FC change cannot be fully explained by lower reliability. Collectively, our results suggest that stable individual differences in baseline FC exert a stronger influence on future cognitive outcomes than changes in FC over time.

### Baseline FC and longitudinal FC change weakly predict cognitive change

We have shown that baseline FC robustly predicted future cognitive ability at Year 2. Longitudinal FC change was also able to predict future cognition at Year 2, though not as well as baseline FC. Here, we further explored whether baseline FC and longitudinal FC change could predict cognitive change between baseline and Year 2. We found that baseline FC could only predict cognitive change for the Rey Auditory Verbal Learning Task (RAVLT; FDR q < 0.05; Figure 6). Similarly, longitudinal FC change could only predict cognitive change of the Little Man Task (LMT; FDR q < 0.05; Figure 6).

**Figure 6.**
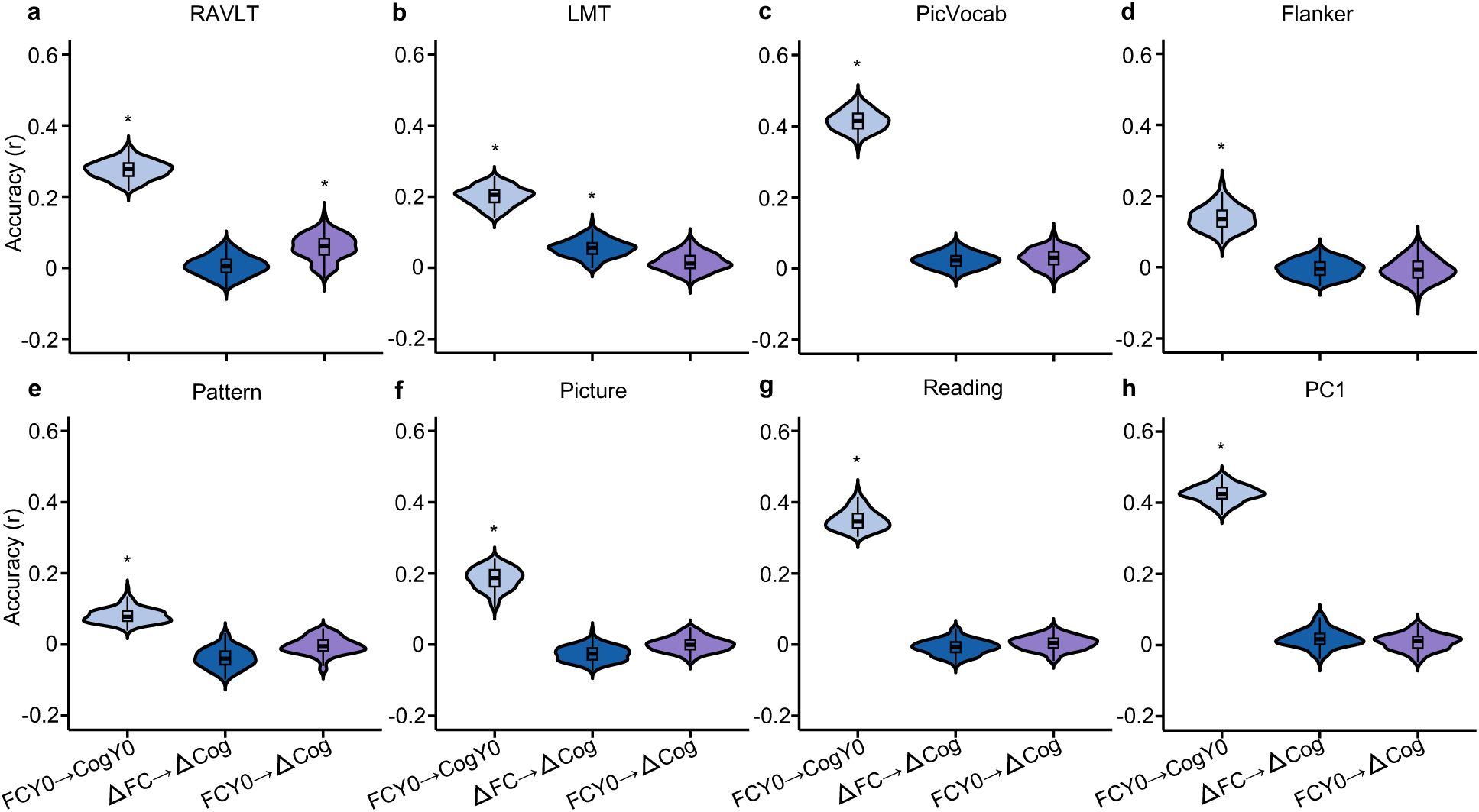
Limited prediction of cognitive change from baseline FC and longitudinal FC change. Each panel corresponds to the prediction accuracy of a different cognitive measure. Within each panel, baseline FC and longitudinal FC change (ΔFC) were used to predict cognitive change (ΔCog). Predictions of baseline cognition from baseline FC (FCY0 → CogY0) were also shown for reference. Asterisks (*) denote above chance prediction after multiple comparisons correction (FDR q < 0.05). RAVLT: Rey Auditory Verbal Learning Test (verbal memory); LMT: Little Man Task (spatial reasoning); PicVocab: Picture Vocabulary Task (vocabulary); Flanker: Flanker Task (executive function); Pattern: Pattern Comparison Processing Speed Test (processing speed); Picture: Picture Sequence Memory Test (episodic memory); Reading: Oral Reading Recognition Task (reading ability). PC1: the first principal component of the above seven cognitive measures.

Importantly, we note that baseline FC did not perform better than longitudinal FC change, suggesting that the weak prediction cannot be fully explained by lower reliability of longitudinal FC change. However, the weak prediction could be due to lower reliability of cognition change compared with baseline cognition. Similar conclusions were reached when using longitudinal rate of changes in FC and cognition between baseline and Year 2 (Figure S7).

### Convergent and divergent predictive network features between cross-sectional and longitudinal estimates of brain–cognition relationship

Consistent with previous studies (Sripada et al., 2020; Chen et al., 2022; Keller et al., 2023; Zhi et al., 2024), we found strong FC–cognition relationship at baseline (Figure 4). For example, individuals with stronger FC within the salience network at baseline also exhibited better cognitive performance at baseline (Figure 4c). Assuming a causal relationship, we might expect individuals with greater increase in salience network FC to exhibit a greater improvement in cognitive performance between the two time points. Since longitudinal FC change only significantly predicted changes in the Little Man Task (LMT; Figure 6b), we focused our analysis on LMT.

More specifically, we computed the predictive network features (PNFs) for the model using baseline FC to predict baseline LMT score (Figure 7a), as well as for the model using longitudinal FC change to predict longitudinal LMT performance change across the two timepoints (Figure 7b). The two PNFs were only weakly correlated (r = 0.22). A spin test excluding subcortical regions (Fischl et al., 2002) confirmed statistical significance (r = 0.17; pSpin < 0.001).

**Figure 7.**
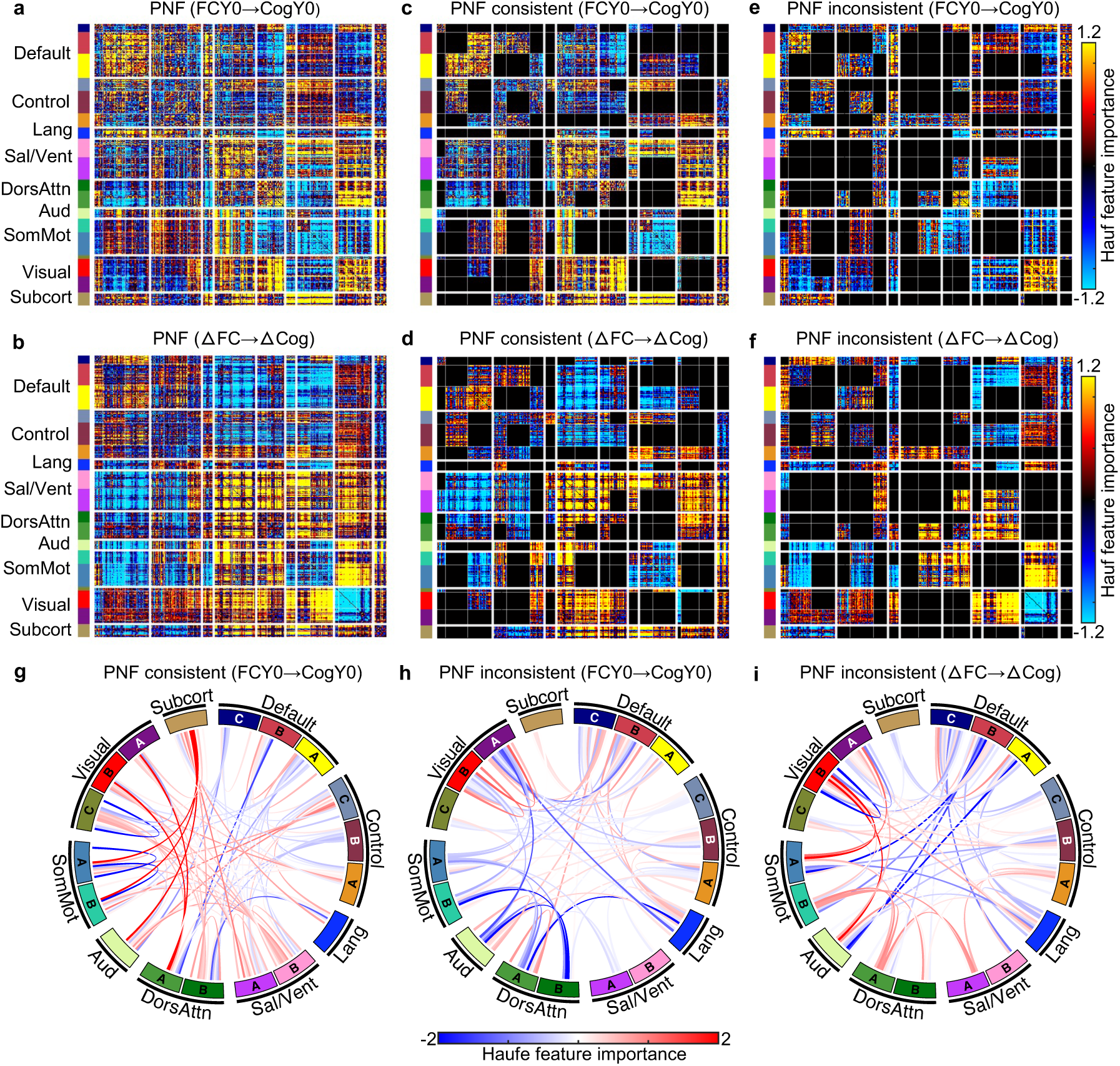
Convergent and divergent predictive network features (PNFs) between cross-sectional and longitudinal estimates of FC–cognition relationship. (a) PNFs from cross-sectional model using baseline FC (FCY0) to predict baseline Little Man Task (LMT) score (CogY0). (b) PNFs from longitudinal model using changes in FC (ΔFC) to predict changes in LMT performance (ΔCog) across the two timepoints. (c) Network blocks with consistent PNFs in the cross-sectional model. (d) Network blocks with consistent PNFs in the longitudinal model. PNFs were considered consistent if the average feature value for a network block had the same sign in both cross-sectional and longitudinal models. (e) Network blocks with inconsistent PNFs in the cross-sectional model. (f) Network blocks with inconsistent PNFs in the longitudinal model. PNFs were considered inconsistent if the average feature value had opposite signs across models. (g) Same as (c), visualized using a chord diagram. (h) Same as (e), visualized using a chord diagram. (i) Same as (f), visualized using a chord diagram. For visualization purposes, each predictive network feature matrix was normalized by dividing all values by the standard deviation of the entire matrix.

Regardless of statistical significance, there were clear convergences (Figures 7c and 7d) and divergences (Figures 7e and 7f) between the cross-sectional and longitudinal PNFs. For example, individuals who exhibited greater increases in FC within the salience network over time also showed larger improvements in LMT performance (Figure 7d). This aligns with the cross-sectional finding that stronger salience network FC at baseline was associated with better LMT performance (Figure 7c), indicating convergence between cross-sectional and longitudinal estimates of brain–cognition relationship (Figure 1b). A chord diagram was used to visualize these convergent links (Figure 7g).

In contrast, individuals with stronger baseline FC within the visual networks tended to perform better on the LMT at baseline (Figures 7e). However, greater longitudinal reduction in visual connectivity predicted larger improvements in LMT performance (Figures 7f), reflecting divergence between cross-sectional and longitudinal estimates of brain–cognition relationship (Figure 1c). Chord diagrams were used to highlight the divergent links across models (Figures 7h and 7i).

For completeness, PNFs for the other seven cognitive measures were presented in Figure S8. Correlations between cross-sectional and longitudinal models of brain–cognition relationship ranged from-0.08 to 0.62. Consistent with LMT, there were clear convergences and divergences between cross-sectional and longitudinal PNFs. However, although the cross-sectional models predicted the seven cognitive measures better than chance, this was not the case for the longitudinal models. As such, the longitudinal PNFs should be interpreted with care.

## DISCUSSION

Using longitudinal rs-fMRI and cognitive measures collected at baseline and Year 2 of the ABCD Study, our findings demonstrate that baseline FC predicts future cognitive ability better than baseline cognitive ability. Furthermore, models trained on baseline FC to predict baseline cognition also improve in accuracy when applied to predict Year 2 cognition from Year 2 FC, suggesting that brain–cognition relationships strengthen over time. Intriguingly, baseline FC emerges as a stronger predictor of future cognition than longitudinal FC change, which cannot be explained by reliability alone. Both baseline FC and longitudinal FC change show limited ability to predict cognitive change. Finally, we observe both convergent and divergent predictive network features between cross-sectional and longitudinal estimates of brain–cognition relationship, revealing a multivariate twist on Simpson’s paradox (Figures 1b and 1c). Together, these findings indicate that although brain–cognition relationships are refined over development, they are primarily shaped by stable individual differences in brain network organization.

### Sustained early individual differences despite longitudinal change in cognition and FC

Developmental science seeks to uncover both stability and change in individual characteristics over time (Emmerich, 1964; Bornstein & Sigman, 1986; Bornstein, Putnick, & Esposito, 2017). In our study, a high similarity in cognitive profiles across two timepoints (Figure 2a) alongside the longitudinal increase in cognitive performance (Figure 2b) reflect developmental stability and change respectively. The brain functional network architecture, as measured by resting-state FC, also exhibited strong developmental stability (Figure 3c) and change (Figure 3e).

Intriguingly, FC stability and change showed regional variations along the sensorimotor-association (S-A) axis (Figure 3d and 3f). The S-A axis represents a key axis of hierarchical cortical organization, spanning from primary sensorimotor to transmodal association cortices (Margulies et al., 2016; Sydnor et al., 2021). Many cross-sectional studies have shown that the development of numerous functional and structural brain properties is organized along the S-A axis (Baum et al., 2022; Larsen et al., 2022; Sydnor et al., 2023; Luo et al., 2024; Zhang et al., 2024). The current study extends these previous studies by providing longitudinal evidence of greater change (and lower stability) in sensory-motor regions and smaller change (and greater stability) in association cortex.

Previous studies have also suggested that association regions mature later than sensorimotor regions (Supekar, Musen, & Menon, 2009; Dosenbach et al., 2010; Cao et al., 2014; Cao et al., 2016; Luo et al., 2024; Sun et al., 2025). The slower maturation of the association cortex might explain our finding that association regions exhibit less longitudinal change than sensorimotor regions across the two ABCD Study timepoints. Specifically, during the developmental period under study, sensorimotor regions may be undergoing relatively rapid development, while the association cortex progresses at a slower pace. At future ABCD Study timepoints, sensorimotor regions may reach a maturation plateau, allowing for more pronounced changes to emerge in the association cortex.

Beyond group-level stability and changes, we also observed substantial individual differences in cognitive (Figure 2c) and functional network (Figure 3g) changes. Intriguingly, there were some participants whose patterns of cognitive improvements (Figure 2d) and FC changes (Figures 3i to 3k) were uncorrelated with other participants. Previous studies have found that sensorimotor and visual regions exhibit lower inter-individual FC variability than association cortex (Mueller et al., 2013; Laumann et al., 2015; Xu et al., 2019; Stoecklein et al., 2020). In contrast, we found that inter-individual differences in FC development were especially pronounced in sensorimotor and visual cortex (Figure 3h). However, the regional variation did not exactly reflect the S-A axis because certain regions of the default network, such as the medial prefrontal cortex also exhibit high variability in longitudinal FC change (Figure 3h). Therefore, our results reveal divergences between inter-individual variation in cross-sectional FC and inter-individual variation in longitudinal FC change.

### Strengthened relationship between FC and cognition during development

Previous studies have shown that brain functional connectomes can reliably predict current and future cognitive performance in children, adolescents, and adults (Rosenberg et al., 2016; Cui et al., 2020; Chen et al., 2022; Ooi et al., 2022; Keller et al., 2023; Zhi et al., 2024). We extend previous findings by demonstrating that baseline FC more accurately predicts future cognitive performance than current cognition, and models of brain–cognition relationship at baseline generalize better when applied to data two years later (Figure 4). The results hold when using a subset of participants with comparable head motion (Figure S4). Thus, our study extends prior understanding by revealing that the brain–cognition relationships strengthen during the transition from childhood to adolescence. These results suggest that functional brain architecture may serve as a stable scaffold that increasingly supports cognitive abilities over time.

### Dominant role of stable individual differences in predicting future cognitive outcomes

A recent study linked individual variability in longitudinal FC change with demographic and maturational factors (Bottenhorn et al., 2023). We build on this work by examining how baseline FC and longitudinal FC changes relate to future cognitive ability. Two prior studies in infancy had divergent findings: one study reported stronger cross-sectional associations (Liu et al., 2021), while another study found stronger longitudinal effects (Salzwedel et al., 2019). These discrepancies may reflect differences in study design, including small sample sizes that can inflate effect sizes (Marek et al., 2022), as well as variation in target region of interest (e.g., amygdala vs. hippocampus) and reliance on univariate analyses. By leveraging a large sample and multivariate prediction methods, our study provides definitive evidence on the relative roles of cross-sectional and longitudinal FC in predicting cognitive outcomes.

Our findings showed that longitudinal change in FC between baseline and Year 2 can predict cognitive performance at Year 2. However, baseline FC outperformed longitudinal FC change in predicting Year 2 cognition (Figures 5a and b). Given that estimates of longitudinal change are often less reliable than cross-sectional estimates due to accumulated measurement noise (Rogosa & Willett, 1983; Noble, Scheinost, & Constable, 2021; Brandmaier, Lindenberger, & McCormick, 2024; Parsons & McCormick, 2024), we developed a method to estimate the reliability of FC change and baseline FC. Even after controlling for reliability, baseline FC still yielded higher predictive accuracy. This suggests that the superior predictive power of baseline FC cannot be fully attributed to reliability differences. Instead, it points to the greater influence of stable individual differences in FC on future cognitive outcomes.

Prior work has shown that individual differences in functional connectome organization are already evident in the third trimester and closely resemble adult-like patterns (Xu et al., 2019; Stoecklein et al., 2020). These early-emerging features, likely shaped by neurogenesis and genetic programming, tend to be relatively stable and less susceptible to postnatal environmental influences (Gao et al., 2014; Xu et al., 2019). In contrast, longitudinal FC change during the transition from childhood and adolescence may reflect more transient, state-dependent dynamics shaped by environmental variability and experience. This contrast may account for the stronger predictive power of baseline FC compared to FC change in forecasting future cognitive outcomes.

Our study also provides estimates of the reliability of longitudinal FC change. Based on 20-minute scans, longitudinal FC change exhibits an ICC of 0.24, while baseline FC exhibits an ICC of 0.56. Because the reliability of longitudinal FC change is determined by the reliabilities of the baseline FC and Year 2 FC, our study suggests that the need for even longer scan duration in longitudinal studies to achieve more reliable estimates of longitudinal FC change (cf. Ooi et al., 2024).

### Convergent and divergent brain–cognition relationships across cross-sectional and longitudinal models

Surprisingly, longitudinal FC changes showed limited ability to predict cognitive change, with only the Little Man Task reaching statistical significance (Figure 6). This cannot be fully explained by the low reliability of longitudinal FC change, given that baseline FC is more reliable but still fails to predict cognitive growth. One possibility is that both FC and its longitudinal changes may be relatively insensitive to the dynamic neurocognitive processes that drive cognitive growth during this transitional period. Furthermore, cognitive development may be strongly shaped by external influences, such as environmental conditions, learning opportunities, and social experiences, which may not be fully captured by FC measures.

However, another possibility is that longitudinal cognition changes are less reliable than baseline cognition, which will lead to worse prediction performance (Nikolaidis et al., 2022; Gell et al., 2024). Indeed, the formula relating cross-sectional FC and longitudinal FC change (Equation 4 in Methods) is also applicable to cognition. The strong correlation of cognitive measures between baseline and Year 2 (Figure 2a) will greatly reduce the reliability of cognitive change. Because of this potential “confound”, we should not over-interpret prediction accuracy differences between cross-sectional (FCY0 → CogY0) and longitudinal (ΔFC → ΔCog) models of brain– cognition relationship (Figure 6). However, we could interpret the Haufe-transformed predictive network features of cognitive measures that were successfully predicted (Figure 7).

We observed both convergence and divergence between cross-sectional and longitudinal predictive network features, extending the classical Simpson’s paradox (Pearl, 2014) (Figure 1a) into a multidimensional developmental context. Stronger baseline salience FC was associated with better baseline LMT performance, and greater increases in salience connectivity over time predicted greater LMT improvement, reflecting convergence between cross-sectional and longitudinal predictive models (Figure 1b). In contrast, while stronger baseline visual FC was associated with better baseline LMT performance, greater reductions in visual connectivity over time predicted greater LMT improvement, indicating divergence between the two estimates (Figure 1c). These findings provide an empirical demonstration of how cross-sectional and longitudinal brain–cognition relationships can converge and diverge, underscoring the need to disentangle their contributions to cognitive development.

### Limitations and future work

At the time of this study, the third timepoint of the ABCD Study had not yet been fully released. As a result, our analysis focused on the two initial timepoints. While using only two timepoints limits our ability to model nonlinear or individual-specific developmental trajectories of functional connectivity (FC) and cognition (Foulkes & Blakemore, 2018; Rosenberg, Casey, & Holmes, 2018), our study still represents the largest longitudinal dataset currently available for examining brain–cognition relationships during the transition from childhood to early adolescence. Although estimates of FC change based on two timepoints may be noisier than those derived from denser longitudinal sampling, our study provides a valuable early benchmark and testbed for predictive modeling of developmental change. As future waves of ABCD Study and other large-scale studies become available, incorporating more timepoints will enable more fine-grained tracking of individual developmental trajectories and their links to cognitive outcomes. Similarly, expanding analyses to span a broader age range may help uncover developmental patterns that extend beyond early adolescence.

## CONCLUSION

Our large-scale longitudinal analysis reveals that although functional connectome and cognition co-evolve during the transition from childhood to early adolescence, stable individual differences exert a stronger influence on future cognitive ability than short-term neural changes.

## METHODS

### Participants

We considered individuals with rs-fMRI and cognition tests from the ABCD Study across 21 sites. Ethical approval was granted by the Institutional Review Board (IRB) at the University of California, San Diego, as well as by the IRBs of each participating study site. Written informed consent was obtained from parents and guardians. At the start of this study, the third time point of ABCD imaging data was still in its initial stage. Thus, we included only participants with rs-fMRI and cognition data at baseline and Year 2, ensuring that all participants were unrelated and remained at the same site across the two time points. Specifically, after rs-fMRI quality control of both time points, 4615 participants remained. Excluding participants lacking cognition measures at both time points reduced the sample size to 3455, and further restricting to unrelated participants resulted in 3147 participants. Finally, we excluded participants who were scanned at different sites across the two time points and those from sites with fewer than 10 participants.

Our final sample comprised 2949 individuals (Table 1).

### Imaging data preprocessing

Minimally processed T1 and rs-fMRI data were utilized. Details about acquisition protocol and minimal processing can be found elsewhere (Casey et al., 2018; Hagler et al., 2019). Resting-state fMRI data preprocessing was in line with our previous study (Chen et al., 2022; Ooi et al., 2022). Specifically, (1) rs-fMRI data were aligned to T1 images using boundary-based registration (Greve & Fischl, 2009). (2) Respiratory pseudo-motion was filtered by applying a band-stop criteria of 0.31–0.43 Hz (Fair et al., 2020). (3) Volumes with framewise displacement (FD) > 0.3 mm or voxel-wise differentiated signal variance (DVARS) > 50 were flagged. Then, each flagged frame, along with the one immediately before and the two immediately after, was censored. Uncensored data segments with fewer than five frames were also censored. (4) Global signal, white matter signal, and ventricular signals, six motion parameters as well as their temporal derivatives were regressed out from the rs-fMRI data, with regression coefficients estimated from uncensored data. (5) The Lomb-Scargle periodogram method (Power et al., 2014) was used to interpolate censored frames. (6) A band-pass filter (0.009–0.08 Hz) was applied. (7) The data were mapped onto fsaverage6 surface space in FreeSurfer and then smoothed with a 6 mm full width at half-maximum kernel.

### Functional connectivity

We constructed the FC matrix by combining the 400-region Yan parcellation (Yan et al., 2023) (Figure 3a) with 19 subcortical regions of interest (ROIs, Figure 3b) defined by Fischl et al. (Fischl et al., 2002). FC was calculated as the Pearson correlation coefficients between the average time series of each ROI pair, resulting in a 419 × 419 FC matrix. Notably, censored frames were excluded when computing FC. We averaged the FC matrices across runs for each participant after Fisher’s r-to-z transformation and converted back to r values after averaging.

### Cognition assessment

We considered seven cognitive tasks administered at both timepoints. Five were drawn from the NIH Toolbox (i.e., Picture Vocabulary Task, Flanker Task, Pattern Comparison Processing Speed Test, Picture Sequence Memory Test, and the Oral Reading Recognition Task) (Weintraub et al., 2013), and two additional tasks included the Rey Auditory Verbal Learning Test (RAVLT) and Little Man Task (LMT) (Luciana et al., 2018).The Picture Vocabulary Task assesses language skills and verbal intelligence, while the Oral Reading Recognition Task measures reading ability by asking participants to pronounce isolated words. The Pattern Comparison Processing Speed Test evaluates rapid visual processing. The Picture Sequence Memory Test assesses episodic memory through the recall of image sequences. The Flanker Task measures response inhibition and conflict monitoring by requiring participants to modulate responses under congruent versus incongruent conditions. The RAVLT assesses auditory learning, memory, and recognition, whereas the LMT engages visual–spatial processing, specifically mental rotation, across varying levels of difficulty.

Consistent with previous work (Thompson et al., 2019), we used uncorrected standard scores for each NIH Toolbox task, total correct scores for the RAVLT, and percent correct scores for the LMT in our analyses.

To obtain a measure of overall cognitive ability, we z-normalized the seven cognitive measures and then applied principal component analysis to derive the first principal component (PC1) explaining the most variability in cognition across participants. To avoid data leakage, the cognitive component was estimated from 6050 participants at baseline that were not included in the main analysis. The loadings of PC1 were then transferred to derive PC1 for the 2949 participants at baseline and Year 2. We note that the z-normalization for the 2949 participants was performed using the mean and standard deviation calculated from the 6050 participants.

### Longitudinal cognitive change

We first applied longitudinal ComBat (Beer et al., 2020) to remove site effects from the cognitive scores. Baseline age, age interval between the two timepoints, and sex were included in the ComBat model to preserve biologically meaningful variability. The output consisted of harmonized baseline and Year 2 cognitive scores, with both additive and multiplicative site effects accounted for.

To examine the stability of cognitive performance between baseline and Year 2, for each of the eight cognitive measures, we computed the Spearman’s correlation of the cognitive measures between the two time points (across participants; Figure 2a). More specifically, we regressed baseline age and sex from the harmonized baseline cognitive scores, as well as Year 2 age and sex from the harmonized Year 2 cognition. Spearman’s correlation was then computed using the resulting residuals.

To examine the longitudinal change in cognitive performance (Figure 2b), we employed a linear mixed effect model consistent with the longitudinal ComBat model. More specifically, baseline age, age interval between the two timepoints and sex are used as covariates.

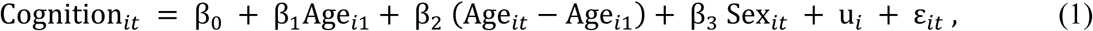

where *i* indexes the *i*-th participant, *t* references the time point, u*i* is the random intercept and ε*it* is the noise term. The t statistic for the age interval coefficient (β2) was subsequently converted to Cohen’s d to quantify the longitudinal effect size. Cognition*it* is the cognition of the *i*-th participant at timepoint *t* after longitudinal ComBat.

However, we note that Equation 1 can only capture a group-level estimate of longitudinal change. Therefore, to capture the potential individual differences in longitudinal cognition change, for each participant and each cognitive score, we computed the longitudinal cognition change as: Cognition*i*2 - Cognition*i*1, where Cognition*it* is the cognition of the *i*-th participant at timepoint *t* (from longitudinal ComBat). We then regressed out sex, baseline age and age interval (i.e., between Year 2 and baseline) from the individual-level longitudinal cognition change estimate.

### Longitudinal functional brain network change

We first applied longitudinal ComBat (Beer et al., 2020) to remove site effects from each functional connectivity (FC) entry. Baseline age, age interval between the two timepoints, sex and head motion were included in the ComBat model to preserve biologically meaningful variability. The output consisted of harmonized baseline and Year 2 functional connectivity (FC) with additive and multiplicative site effects accounted for.

To examine the stability of FC between baseline and Year 2, for each FC entry, we computed the Spearman’s correlation of the FC value between the two time points (across participants; Figure 3c). More specifically, we regressed baseline age, sex and mean head motion (as measured by framewise displacement FD) from the harmonized baseline FC, as well as Year 2 age, sex and mean head motion (FD) from the harmonized Year 2 FC. Spearman’s correlation was then computed using the resulting residuals.

To examine the longitudinal change in FC (Figure 3e), we employed a linear mixed effect model consistent with the longitudinal ComBat model. More specifically, baseline age, age interval between the two timepoints, sex, and mean FD were included as covariates.

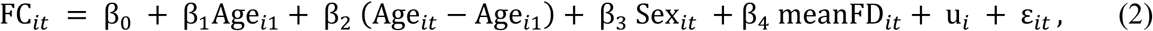

where *i* indexes the *i*-th participant, *t* references the time point, u*i* is the random intercept and ε*it* is the noise term. The t statistic for the age interval coefficient (β2) was subsequently converted to Cohen’s d to quantify the longitudinal effect size. FC*it* is the FC value of a particular FC edge of the *i*-th participant at timepoint *t* after longitudinal ComBat.

However, we note that Equation 2 captures only the group-level estimate of longitudinal change. Therefore, to capture the potential individual differences in longitudinal FC change, for each participant and each FC edge, we computed the z value of FC change across the two timepoints:

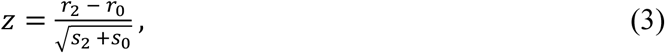

where *r*2 and *r*0 denote Fisher r-to-z transformed FC value for the FC edge at Year 2 and baseline respectively. *s*2 and *s*0 denote the variance of the FC value for each FC edge at Year 2 and baseline respectively. The estimated variance needed to account for auto-correlation in the fMRI time series, so we used the MATLAB function xDF.m (https://github.com/asoroosh/xDF) to estimate *r* and *s* for each FC edge at each time point (Afyouni, Smith, & Nichols, 2019). We then regressed sex, baseline age and age interval (between Year 2 and baseline), mean head motion (FD) at baseline, mean head motion (FD) at Year 2 from the z-statistic across participants.

### Using cross-sectional FC to predict cross-sectional cognition

To explore the cross-sectional relationship between FC and cognition, we used (1) baseline FC to predict baseline cognition, and also (2) Year 2 FC to predict Year 2 cognition (Figures 4a & 4b). For comparison, we also used baseline FC to predict Year 2 cognition.

For the prediction analysis, we applied the same KRR framework as in our previous studies (Chen et al., 2022; Ooi et al., 2022). Briefly, participants from all imaging sites were grouped into 10 “site clusters”, with each cluster comprising all participants from one or more sites and containing at least 280 individuals.

We then implemented a leave-3-site-cluster-out nested cross-validation approach, where 7 random site clusters were used as the training set, and the remaining 3 clusters served as the test set. This process resulted in 120 unique replications, covering every possible split. We emphasize that individuals from the same site were not split across site clusters, so the test set always contains participants not from the same site as the training set.

10-fold cross-validation was performed within the training set to determine the optimal regularization hyperparameter. The best hyperparameter was used to train a final model from the full training set. This final model was then applied to the test set. This procedure was repeated 120 times, covering every possible split.

When using baseline FC to predict baseline cognition, we controlled for sex, baseline age, and baseline head motion. When using Year 2 FC to predict Year 2 cognition, we controlled for sex, Year 2 age, and Year 2 head motion (mean FD). When using baseline FC to predict Year 2 cognition, we controlled for sex, Year 2 age, age interval between the two timepoints, and baseline head motion (mean FD). All regressions were performed on the training set, and the resulting regression coefficients were applied to the test set.

Prediction accuracy was calculated as the Pearson’s correlation between the predicted scores and actual scores within each test set and then averaged across test sets. We note that in this prediction analysis (and all future prediction analyses), longitudinal ComBat was not performed because ComBat requires FC and/or cognition from all sites (including those from the test set) to be included in the mixed effects model, resulting in test set leakage.

Not performing any harmonization would in theory hurt our prediction accuracy, as opposed to inflating our prediction accuracy. Furthermore, by performing leave-3-site-cluster-out cross-validation, our prediction procedure is generalizable to hypothetical new sites in which there is only a single individual, so harmonization cannot be performed. Therefore, we believe that our approach is a reasonable (i.e., conservative) course of action.

### Model interpretation

To interpret feature importance in the predictive models (Figure 4c), we employed the Haufe transformation to yield a 419 × 419 predictive network feature (PNF) matrix for each cognitive measure (Haufe et al., 2014; Chen et al., 2022; Ooi et al., 2022; Chen et al., 2023). A positive PNF indicates that a higher FC value was associated with a higher predicted cognitive score.

### Model transfer across time points

We have previously trained a model to use baseline FC to predict baseline cognition. Here, we examined whether the model can be used to predict cognition at Year 2 (Figure 4d) without any further tuning.

More specifically, for a given split of the 10 site clusters into a training set (7 site clusters) and a test set (3 site clusters), we trained the baseline model on the baseline training set, after regressing out sex, baseline age and baseline head motion from baseline cognition. The regression coefficients for sex, age and head motion (from the training set) were then applied to regress sex, Year 2 age and Year 2 head motion from Year 2 cognition in the test set. Finally, we used the baseline model to predict Year 2 cognition (after regressing out the covariates) from Year 2 FC.

To control for the influence of head motion on our results, we selected a subset of participants (n = 2642) with matched mean FD across the two time points and repeated the entire model transfer analyses (Figure S4).

### Using longitudinal FC change (delta) to predict Year 2 cognition

We explored the use of FC change (delta) to predict cognition at Year 2 using KRR (Figures 5a & 5b). FC change (or delta) is defined as the difference between Year 2 FC and baseline FC. Here, we controlled for sex, baseline age, age interval, baseline head motion (mean FD) and Year 2 head motion (mean FD). All regressions were performed on the training set, and the resulting regression coefficients were applied to the test set.

As a control analysis, we repeated the analyses using rate of FC change (Figure S6). Rate of FC change was defined as (FCY2-FCY0)/(AgeY2-AgeY0). The same set of nuisance regressors was used.

### Mathematical relationship between the reliability of FC change and reliability of cross-sectional FC

As shown in the main results, longitudinal FC change was less effective than baseline FC in predicting Year 2 cognition. One contributing factor could be the lower reliability of FC change, compared with cross-sectional baseline FC. To illustrate this, we derived the mathematical relationship between the reliability of FC change and that of cross-sectional FC (Rogosa & Willett, 1983).

Let *R*1 be the reliability of FC at timepoint 1 and *R*2 be the reliability of FC at timepoint 2. Let *V*1 be the variance of FC at timepoint 1 and *V*2 be the variance of FC at timepoint 2. Let *ρ* be the empirical Pearson’s correlation between the FC of the two timepoints. The reliability of the longitudinal FC change, denoted *RD*, can then be expressed as (see Supplementary Methods):

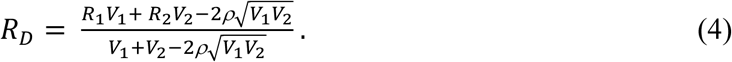

A higher *ρ* will lead to lower FC change reliability (*RD*). Indeed, the high FC stability between baseline and Year 2 (Figures 3c and 3d) implies a large *ρ*, and consequently, a lower FC change reliability (*RD*).

### Empirically estimating the reliability of cross-sectional FC and longitudinal FC change

To investigate whether the lower FC change reliability could explain the weaker prediction, we first empirically estimate the reliability of cross-sectional (baseline) FC and longitudinal FC change. However, there is no longitudinal test-retest data in ABCD Study, so we developed a method to estimate cross-sectional FC reliability and longitudinal FC change reliability.

More specifically, to estimate cross-sectional (baseline) FC reliability, we chose a subset of participants with 4 runs after quality control (n = 897) and divide the 4 runs into 2 groups comprising the first two runs and the last two runs. We can compute FC twice, once using the first two runs (10 minutes) and once using the last two runs (10 minutes). By treating the two FC estimates as test-retest data, we can compute ICC using the one-way random effects ANOVA model (Shrout & Fleiss, 1979):

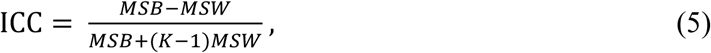

where *MSB* denotes the mean square between-participants variability, *MSW* denotes mean square within-participant variability, and *K* is the number of repeat measures (which is two in this case). However, this ICC estimate (based on 10 minutes of fMRI) is probably smaller than the ICC of FC based on the full 20 min of fMRI.

To estimate the ICC of the baseline FC based on the full 20 minutes of fMRI, we adopt the following approximation (Ooi et al., 2024):

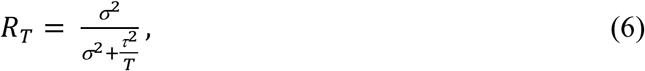

where *RT* is the reliability based on *T* min of fMRI. Τ^2^/*T* represents the variance of the fMRI noise, which decreases with scan time *T*, but the variance is modulated by τ^2^(due to autocorrelation in the fMRI signal). Finally, σ^2^ is the true between-participant variability. As scan time *T* becomes large, the noise component diminishes, and reliability asymptotically approaches 1.

We would like to use Equation 6 to infer the reliability of cross-sectional (baseline) FC based on 20 min of fMRI, but σ^2^ and *τ*^2^ are unknown. Therefore, we computed ICC of cross-sectional FC using 2, 4, 6, 8 and 10 minutes of fMRI. We then applied Equation 6 to estimate σ^2^ and *τ*^2^, and substituted *T* = 20 minutes into Equation 6 to obtain the cross-sectional (baseline) FC reliability based on 20 minutes of fMRI. Note that σ^2^ and *τ*^2^ are generally different for different FC edge, so we repeated the whole procedure independently for all FC edges. The fit of Equation 6 is extremely good in practice, with a median coefficient of determination (COD) of 0.95.

We repeated the same procedure to obtain FC reliability at Year 2 obtaining a strong fit (median COD = 0.94). Finally, we applied Equation 4 to obtain the reliability of longitudinal FC change.

### Controlling for reliability when comparing baseline FC and longitudinal FC change for predicting Year 2 cognition

As shown in the main results, longitudinal FC change was less predictive of Year 2 cognition than baseline FC. To test whether this prediction gap could be attributed to the lower reliability of longitudinal FC change, we reduced the scan duration used to compute baseline FC so that its reliability would match that of longitudinal FC change. Specifically, we identified the value of *T* in Equation 6 that equated baseline FC reliability to longitudinal FC reliability. This value of *T* varied across FC edges, with a median of 4.66 minutes and a mean of 5.05 minutes. To adopt a conservative approach, we fixed *T* at 4 minutes. We then used baseline FC estimated from 4 minutes of fMRI data to predict Year 2 cognition (Figures 5a and 5b).

### Using longitudinal FC change (delta) to predict longitudinal cognitive change

To investigate the divergence and convergence of longitudinal and cross-sectional estimates of brain–cognition relationship, we also used longitudinal FC change to predict the longitudinal cognitive change. We controlled for sex, baseline age, age interval, baseline head motion (mean FD) and Year 2 head motion (mean FD). All regressions were performed on the training set, and the resulting regression coefficients were applied to the test set.

For comparison, we also used baseline FC to predict the cognitive change. Here, we controlled for sex, baseline age, age interval, and baseline head motion (mean FD). Once again, all regressions were performed on the training set, and the resulting regression coefficients were applied to the test set.

As a control analysis, we repeated the analyses using rate of FC change instead of FC change and rate of cognitive change instead of cognitive change (Figure S7). We controlled for sex, baseline age, age interval, baseline head motion (mean FD) and Year 2 head motion (mean FD). All regressions were performed on the training set, and the resulting regression coefficients were applied to the test set.

### Statistical tests of prediction accuracy

To test whether prediction accuracy was better than chance, permutation tests were performed by shuffling cognitive measures across participants 1,000 times within sites and then repeating the leave-3-site-cluster-out cross-validation procedure.

To compare prediction accuracies between models, we employed the corrected resampled t-test (Nadeau & Bengio, 2003), which corrected for dependencies across cross-validation folds.

All p-values were computed based on two tails. Multiple comparisons were controlled using the false discovery rate (FDR) with q < 0.05 (Benjamini & Hochberg, 1995).

### Statistical tests with spatial permutation

To assess the significance of correlations between ICCs from 4-minute baseline FC and 20-minute FC change, as well as between PNFs derived from baseline and longitudinal models, we used a spatial permutation (“spin”) test (Alexander-Bloch et al., 2018; Vasa et al., 2018). A total of 400 cortical parcels (Yan et al., 2023) were randomly rotated on the spherical surface 1000 times to generate a null distribution while preserving spatial autocorrelation. For each permutation, the rows and columns of the matrix were reordered according to the rotated parcellation labels, and a null correlation was computed between the permuted matrix and the original comparison matrix. The observed correlation was then compared against this null distribution to determine significance.

## ACKNOWLEGEMENT

Our research is supported by the NUS Yong Loo Lin School of Medicine (NUHSRO/2020/124/ TMR /LOA), the Singapore National Medical Research Council (NMRC) LCG (OFLCG19May-0035), NMRC CTG-IIT (CTGIIT23jan-0001), NMRC OF-IRG (OFIRG24jan-0006; OFIRG24jul-0049), NMRC STaR (STaR20nov-0003), Singapore Ministry of Health (MOH) Centre Grant (CG21APR1009), and the United States National Institutes of Health (R01MH133334). Any opinions, findings and conclusions or recommendations expressed in this material are those of the authors and do not reflect the views of the funders.

## REFERENCE

Afyouni, S., Smith, S. M., & Nichols, T. E. (2019). Effective degrees of freedom of the Pearson’s correlation coefficient under autocorrelation. Neuroimage, 199, 609–625. 10.1016/j.neuroimage.2019.05.011

Alexander-Bloch, A. F., Shou, H., Liu, S., Satterthwaite, T. D., Glahn, D. C., Shinohara, R. T., Vandekar, S. N., & Raznahan, A. (2018). On testing for spatial correspondence between maps of human brain structure and function. Neuroimage, 178, 540–551. 10.1016/j.neuroimage.2018.05.070

Anokhin, A. P., Luciana, M., Banich, M., Barch, D., Bjork, J. M., Gonzalez, M. R., Gonzalez, R., Haist, F., Jacobus, J., Lisdahl, K., McGlade, E., McCandliss, B., Nagel, B., Nixon, S. J., Tapert, S., Kennedy, J. T., & Thompson, W. (2022). Age-related changes and longitudinal stability of individual differences in ABCD Neurocognition measures. Dev Cogn Neurosci, 54, 101078. 10.1016/j.dcn.2022.101078

Baum, G. L., Flournoy, J. C., Glasser, M. F., Harms, M. P., Mair, P., Sanders, A. F., Barch, D. M., Buckner, R. L., Bookheimer, S., & Dapretto, M. (2022). Graded variation in T1w/T2w ratio during adolescence: measurement, caveats, and implications for development of cortical myelin. Journal of Neuroscience, 42(29), 5681–5694.

Beer, J. C., Tustison, N. J., Cook, P. A., Davatzikos, C., Sheline, Y. I., Shinohara, R. T., Linn, K. A., & Alzheimer’s Disease Neuroimaging, I. (2020). Longitudinal ComBat: A method for harmonizing longitudinal multi-scanner imaging data. Neuroimage, 220, 117129. 10.1016/j.neuroimage.2020.117129

Benjamini, Y., & Hochberg, Y. (1995). Controlling the False Discovery Rate: A Practical and Powerful Approach to Multiple Testing. J R Stat Soc Series B Stat Methodol, 57(1), 289–300. 10.1111/j.2517-6161.1995.tb02031.x

Biswal, B., Yetkin, F. Z., Haughton, V. M., & Hyde, J. S. (1995). Functional connectivity in the motor cortex of resting human brain using echo-planar MRI. Magn Reson Med, 34(4), 537–541. 10.1002/mrm.1910340409

Bornstein, M. H., & Sigman, M. D. (1986). Continuity in mental development from infancy. Child development, 251-274.

Bornstein, M. H., Putnick, D. L., & Esposito, G. (2017). Continuity and Stability in Development. Child Dev Perspect, 11(2), 113–119. 10.1111/cdep.12221

Bottenhorn, K. L., Cardenas-Iniguez, C., Mills, K. L., Laird, A. R., & Herting, M. M. (2023). Profiling intra-and inter-individual differences in brain development across early adolescence. Neuroimage, 279, 120287. 10.1016/j.neuroimage.2023.120287

Brandmaier, A. M., Lindenberger, U., & McCormick, E. M. (2024). Optimal two-time point longitudinal models for estimating individual-level change: Asymptotic insights and practical implications. Dev Cogn Neurosci, 70, 101450. 10.1016/j.dcn.2024.101450

Cao, M., Huang, H., Peng, Y., Dong, Q., & He, Y. (2016). Toward developmental connectomics of the human brain. Frontiers in neuroanatomy, 10, 25.

Cao, M., Wang, J. H., Dai, Z. J., Cao, X. Y., Jiang, L. L., Fan, F. M., Song, X. W., Xia, M. R., Shu, N., Dong, Q., Milham, M. P., Castellanos, F. X., Zuo, X. N., & He, Y. (2014). Topological organization of the human brain functional connectome across the lifespan. Dev Cogn Neurosci, 7, 76–93. 10.1016/j.dcn.2013.11.004

Casey, B. J., Cannonier, T., Conley, M. I., Cohen, A. O., Barch, D. M., Heitzeg, M. M., Soules, M. E., Teslovich, T., Dellarco, D. V., Garavan, H., Orr, C. A., Wager, T. D., Banich, M. T., Speer, N. K., Sutherland, M. T., Riedel, M. C., Dick, A. S., Bjork, J. M., Thomas, K. M.,…Workgroup, A. I. A. (2018). The Adolescent Brain Cognitive Development (ABCD) study: Imaging acquisition across 21 sites. Dev Cogn Neurosci, 32, 43–54. 10.1016/j.dcn.2018.03.001

Chen, J., Ooi, L. Q. R., Tan, T. W. K., Zhang, S., Li, J., Asplund, C. L., Eickhoff, S. B., Bzdok, D., Holmes, A. J., & Yeo, B. T. (2023). Relationship between prediction accuracy and feature importance reliability: An empirical and theoretical study. Neuroimage, 274, 120115.

Chen, J., Tam, A., Kebets, V., Orban, C., Ooi, L. Q. R., Asplund, C. L., Marek, S., Dosenbach, N. U. F., Eickhoff, S. B., Bzdok, D., Holmes, A. J., & Yeo, B. T. T. (2022). Shared and unique brain network features predict cognitive, personality, and mental health scores in the ABCD study. Nat Commun, 13(1), 2217. 10.1038/s41467-022-29766-8

Cui, Z., Li, H., Xia, C. H., Larsen, B., Adebimpe, A., Baum, G. L., Cieslak, M., Gur, R. E., Gur, R. C., Moore, T. M., Oathes, D. J., Alexander-Bloch, A. F., Raznahan, A., Roalf, D. R., Shinohara, R. T., Wolf, D. H., Davatzikos, C., Bassett, D. S., Fair, D. A.,… Satterthwaite, T. D. (2020). Individual Variation in Functional Topography of Association Networks in Youth. Neuron, 106(2), 340–353 e348. 10.1016/j.neuron.2020.01.029

Curran, P. J., & Bauer, D. J. (2011). The disaggregation of within-person and between-person effects in longitudinal models of change. Annu Rev Psychol, 62, 583–619. 10.1146/annurev.psych.093008.100356

Di Biase, M. A., Tian, Y. E., Bethlehem, R. A. I., Seidlitz, J., Alexander-Bloch, A. F., Yeo, B. T. T., & Zalesky, A. (2023). Mapping human brain charts cross-sectionally and longitudinally. Proc Natl Acad Sci U S A, 120(20), e2216798120. 10.1073/pnas.2216798120

Dosenbach, N. U., Nardos, B., Cohen, A. L., Fair, D. A., Power, J. D., Church, J. A., Nelson, S. M., Wig, G. S., Vogel, A. C., Lessov-Schlaggar, C. N., Barnes, K. A., Dubis, J. W., Feczko, E., Coalson, R. S., Pruett, J. R., Jr., Barch, D. M., Petersen, S. E., & Schlaggar, B. L. (2010). Prediction of individual brain maturity using fMRI. Science, 329(5997), 1358–1361. 10.1126/science.1194144

Emmerich, W. (1964). Continuity and stability in early social development. Child development, 311-332.

Fair, D. A., Miranda-Dominguez, O., Snyder, A. Z., Perrone, A., Earl, E. A., Van, A. N., Koller, J. M., Feczko, E., Tisdall, M. D., & van der Kouwe, A. (2020). Correction of respiratory artifacts in MRI head motion estimates. Neuroimage, 208, 116400.

Finn, E. S., Shen, X., Scheinost, D., Rosenberg, M. D., Huang, J., Chun, M. M., Papademetris, X., & Constable, R. T. (2015). Functional connectome fingerprinting: identifying individuals using patterns of brain connectivity. Nat Neurosci, 18(11), 1664–1671. 10.1038/nn.4135

Fischl, B., Salat, D. H., Busa, E., Albert, M., Dieterich, M., Haselgrove, C., Van Der Kouwe, A., Killiany, R., Kennedy, D., & Klaveness, S. (2002). Whole brain segmentation: automated labeling of neuroanatomical structures in the human brain. Neuron, 33(3), 341–355.

Foulkes, L., & Blakemore, S. J. (2018). Studying individual differences in human adolescent brain development. Nat Neurosci, 21(3), 315–323. 10.1038/s41593-018-0078-4

Fox, M. D., Snyder, A. Z., Zacks, J. M., & Raichle, M. E. (2006). Coherent spontaneous activity accounts for trial-to-trial variability in human evoked brain responses. Nat Neurosci, 9(1), 23–25. 10.1038/nn1616

Gao, W., Elton, A., Zhu, H., Alcauter, S., Smith, J. K., Gilmore, J. H., & Lin, W. (2014). Intersubject variability of and genetic effects on the brain’s functional connectivity during infancy. J Neurosci, 34(34), 11288–11296. 10.1523/JNEUROSCI.5072-13.2014

Gell, M., Eickhoff, S. B., Omidvarnia, A., Kuppers, V., Patil, K. R., Satterthwaite, T. D., Muller, V. I., & Langner, R. (2024). How measurement noise limits the accuracy of brain-behaviour predictions. Nat Commun, 15(1), 10678. 10.1038/s41467-024-54022-6

Gopnik, A., O’Grady, S., Lucas, C. G., Griffiths, T. L., Wente, A., Bridgers, S., Aboody, R., Fung, H., & Dahl, R. E. (2017). Changes in cognitive flexibility and hypothesis search across human life history from childhood to adolescence to adulthood. Proc Natl Acad Sci U S A, 114(30), 7892–7899. 10.1073/pnas.1700811114

Greve, D. N., & Fischl, B. (2009). Accurate and robust brain image alignment using boundary-based registration. Neuroimage, 48(1), 63–72.

Guillaume, B., Hua, X., Thompson, P. M., Waldorp, L., Nichols, T. E., & Alzheimer’s Disease Neuroimaging, I. (2014). Fast and accurate modelling of longitudinal and repeated measures neuroimaging data. Neuroimage, 94, 287–302. 10.1016/j.neuroimage.2014.03.029

Hagler, D. J., Jr., Hatton, S., Cornejo, M. D., Makowski, C., Fair, D. A., Dick, A. S., Sutherland, M. T., Casey, B. J., Barch, D. M., Harms, M. P., Watts, R., Bjork, J. M., Garavan, H. P., Hilmer, L., Pung, C. J., Sicat, C. S., Kuperman, J., Bartsch, H., Xue, F.,… Dale, A. M. (2019). Image processing and analysis methods for the Adolescent Brain Cognitive Development Study. Neuroimage, 202, 116091. 10.1016/j.neuroimage.2019.116091

Haufe, S., Meinecke, F., Görgen, K., Dähne, S., Haynes, J.-D., Blankertz, B., & Bießmann, F. (2014). On the interpretation of weight vectors of linear models in multivariate neuroimaging. Neuroimage, 87, 96–110.

Jernigan, T. L., Brown, S. A., & Dowling, G. J. (2018). The adolescent brain cognitive development study. Journal of research on adolescence: the official journal of the Society for Research on Adolescence, 28(1), 154.

Kang, K., Seidlitz, J., Bethlehem, R. A. I., Xiong, J., Jones, M. T., Mehta, K., Keller, A. S., Tao, R., Randolph, A., Larsen, B., Tervo-Clemmens, B., Feczko, E., Dominguez, O. M., Nelson, S. M., Lifespan Brain Chart, C., Schildcrout, J., Fair, D. A., Satterthwaite, T. D., Alexander-Bloch, A., & Vandekar, S. (2024). Study design features increase replicability in brain-wide association studies. Nature. 10.1038/s41586-024-08260-9

Keller, A. S., Pines, A. R., Shanmugan, S., Sydnor, V. J., Cui, Z., Bertolero, M. A., Barzilay, R., Alexander-Bloch, A. F., Byington, N., Chen, A., Conan, G. M., Davatzikos, C., Feczko, E., Hendrickson, T. J., Houghton, A., Larsen, B., Li, H., Miranda-Dominguez, O., Roalf, D. R.,… Satterthwaite, T. D. (2023). Personalized functional brain network topography is associated with individual differences in youth cognition. Nat Commun, 14(1), 8411. 10.1038/s41467-023-44087-0

Kong, R., Li, J., Orban, C., Sabuncu, M. R., Liu, H., Schaefer, A., Sun, N., Zuo, X. N., Holmes, A. J., Eickhoff, S. B., & Yeo, B. T. T. (2019). Spatial Topography of Individual-Specific Cortical Networks Predicts Human Cognition, Personality, and Emotion. Cereb Cortex, 29(6), 2533–2551. 10.1093/cercor/bhy123

Kong, R., Yang, Q., Gordon, E., Xue, A., Yan, X., Orban, C., Zuo, X. N., Spreng, N., Ge, T., Holmes, A., Eickhoff, S., & Yeo, B. T. T. (2021). Individual-Specific Areal-Level Parcellations Improve Functional Connectivity Prediction of Behavior. Cereb Cortex, 31(10), 4477–4500. 10.1093/cercor/bhab101

Larsen, B., Cui, Z., Adebimpe, A., Pines, A., Alexander-Bloch, A., Bertolero, M., Calkins, M. E., Gur, R. E., Gur, R. C., & Mahadevan, A. S. (2022). A developmental reduction of the excitation: inhibition ratio in association cortex during adolescence. Science advances, 8(5), eabj8750.

Laumann, T. O., Gordon, E. M., Adeyemo, B., Snyder, A. Z., Joo, S. J., Chen, M. Y., Gilmore, A. W., McDermott, K. B., Nelson, S. M., Dosenbach, N. U., Schlaggar, B. L., Mumford, J. A., Poldrack, R. A., & Petersen, S. E. (2015). Functional System and Areal Organization of a Highly Sampled Individual Human Brain. Neuron, 87(3), 657–670. 10.1016/j.neuron.2015.06.037

Liu, J., Chen, Y., Stephens, R., Cornea, E., Goldman, B., Gilmore, J. H., & Gao, W. (2021). Hippocampal functional connectivity development during the first two years indexes 4-year working memory performance. Cortex, 138, 165–177. 10.1016/j.cortex.2021.02.005

Luciana, M., Bjork, J. M., Nagel, B. J., Barch, D. M., Gonzalez, R., Nixon, S. J., & Banich, M. T. (2018). Adolescent neurocognitive development and impacts of substance use: Overview of the adolescent brain cognitive development (ABCD) baseline neurocognition battery. Dev Cogn Neurosci, 32, 67–79. 10.1016/j.dcn.2018.02.006

Luna, B., Garver, K. E., Urban, T. A., Lazar, N. A., & Sweeney, J. A. (2004). Maturation of cognitive processes from late childhood to adulthood. Child development, 75(5), 1357–1372.

Luo, A. C., Sydnor, V. J., Pines, A., Larsen, B., Alexander-Bloch, A. F., Cieslak, M., Covitz, S., Chen, A. A., Esper, N. B., Feczko, E., Franco, A. R., Gur, R. E., Gur, R. C., Houghton, A., Hu, F., Keller, A. S., Kiar, G., Mehta, K., Salum, G. A.,… Satterthwaite, T. D. (2024). Functional connectivity development along the sensorimotor-association axis enhances the cortical hierarchy. Nat Commun, 15(1), 3511. 10.1038/s41467-024-47748-w

Marek, S., Tervo-Clemmens, B., Calabro, F. J., Montez, D. F., Kay, B. P., Hatoum, A. S., Donohue, M. R., Foran, W., Miller, R. L., Hendrickson, T. J., Malone, S. M., Kandala, S., Feczko, E., Miranda-Dominguez, O., Graham, A. M., Earl, E. A., Perrone, A. J., Cordova, M., Doyle, O.,… Dosenbach, N. U. F. (2022). Reproducible brain-wide association studies require thousands of individuals. Nature, 603(7902), 654–660. 10.1038/s41586-022-04492-9

Margulies, D. S., Ghosh, S. S., Goulas, A., Falkiewicz, M., Huntenburg, J. M., Langs, G., Bezgin, G., Eickhoff, S. B., Castellanos, F. X., Petrides, M., Jefferies, E., & Smallwood, J. (2016). Situating the default-mode network along a principal gradient of macroscale cortical organization. Proc Natl Acad Sci U S A, 113(44), 12574–12579. 10.1073/pnas.1608282113

Mueller, S., Wang, D., Fox, M. D., Yeo, B. T., Sepulcre, J., Sabuncu, M. R., Shafee, R., Lu, J., & Liu, H. (2013). Individual variability in functional connectivity architecture of the human brain. Neuron, 77(3), 586–595. 10.1016/j.neuron.2012.12.028

Nadeau, C., & Bengio, Y. (2003). Inference for the Generalization Error. Machine Learning, 52(3), 239–281. 10.1023/a:1024068626366

Nikolaidis, A., Chen, A. A., He, X., Shinohara, R., Vogelstein, J., Milham, M., & Shou, H. (2022). 10.1101/2022.07.22.501193

Noble, S., Scheinost, D., & Constable, R. T. (2021). A guide to the measurement and interpretation of fMRI test-retest reliability. Curr Opin Behav Sci, 40, 27–32. 10.1016/j.cobeha.2020.12.012

Ooi, L. Q. R., Chen, J., Zhang, S., Kong, R., Tam, A., Li, J., Dhamala, E., Zhou, J. H., Holmes, A. J., & Yeo, B. T. T. (2022). Comparison of individualized behavioral predictions across anatomical, diffusion and functional connectivity MRI. Neuroimage, 263, 119636. 10.1016/j.neuroimage.2022.119636

Ooi, L. Q. R., Orban, C., Zhang, S., Nichols, T. E., Tan, T. W. K., Kong, R., Marek, S., Dosenbach, N. U. F., Laumann, T., Gordon, E. M., Yap, K. H., Ji, F., Chong, J. S. X., Chen, C., An, L., Franzmeier, N., Roemer, S. N., Hu, Q., Ren, J.,… Alzheimer’s Disease Neuroimaging, I. (2024). MRI economics: Balancing sample size and scan duration in brain wide association studies. bioRxiv. 10.1101/2024.02.16.580448

Parsons, S., & McCormick, E. M. (2024). Limitations of two time point data for understanding individual differences in longitudinal modeling - What can difference reveal about change? Dev Cogn Neurosci, 66, 101353. 10.1016/j.dcn.2024.101353

Pearl, J. (2014). Understanding Simpson’s Paradox. The American Statistician, 68(1), 8–13. 10.1080/00031305.2014.876829

Piaget, J. (1952). The origins of intelligence in children. International University.

Power, J. D., Mitra, A., Laumann, T. O., Snyder, A. Z., Schlaggar, B. L., & Petersen, S. E. (2014). Methods to detect, characterize, and remove motion artifact in resting state fMRI. Neuroimage, 84, 320–341. 10.1016/j.neuroimage.2013.08.048

Rogosa, D. R., & Willett, J. B. (1983). Demonstrating the Reliability of the Difference Score in the Measurement of Change. Journal of Educational Measurement, 20, 335–343. https://www.jstor.org/stable/1434950

Rosenberg, M. D., Casey, B. J., & Holmes, A. J. (2018). Prediction complements explanation in understanding the developing brain. Nat Commun, 9(1), 589. 10.1038/s41467-018-02887-9

Rosenberg, M. D., Finn, E. S., Scheinost, D., Papademetris, X., Shen, X., Constable, R. T., & Chun, M. M. (2016). A neuromarker of sustained attention from whole-brain functional connectivity. Nat Neurosci, 19(1), 165–171. 10.1038/nn.4179

Saha, R., Saha, D. K., Rahaman, M. A., Fu, Z., Liu, J., & Calhoun, V. D. (2024). A Method to Estimate Longitudinal Change Patterns in Functional Network Connectivity of the Developing Brain Relevant to Psychiatric Problems, Cognition, and Age. Brain Connect, 14(2), 130–140. 10.1089/brain.2023.0040

Salzwedel, A. P., Stephens, R. L., Goldman, B. D., Lin, W., Gilmore, J. H., & Gao, W. (2019). Development of Amygdala Functional Connectivity During Infancy and Its Relationship With 4-Year Behavioral Outcomes. Biol Psychiatry Cogn Neurosci Neuroimaging, 4(1), 62–71. 10.1016/j.bpsc.2018.08.010

Schaefer, A., Kong, R., Gordon, E. M., Laumann, T. O., Zuo, X. N., Holmes, A. J., Eickhoff, S. B., & Yeo, B. T. T. (2018). Local-Global Parcellation of the Human Cerebral Cortex from Intrinsic Functional Connectivity MRI. Cereb Cortex, 28(9), 3095–3114. 10.1093/cercor/bhx179

Shaffer. (2013). Developmental Psychology.

Shrout, P., & Fleiss, J. (1979). Intraclass correlations: Uses in assessing rater reliability. Psychological bulletin, 86, 420–428. 10.1037/0033-2909.86.2.420

Smith, S. M., Fox, P. T., Miller, K. L., Glahn, D. C., Fox, P. M., Mackay, C. E., Filippini, N., Watkins, K. E., Toro, R., Laird, A. R., & Beckmann, C. F. (2009). Correspondence of the brain’s functional architecture during activation and rest. Proc Natl Acad Sci U S A, 106(31), 13040–13045. 10.1073/pnas.0905267106

Sorensen, O., Walhovd, K. B., & Fjell, A. M. (2021). A recipe for accurate estimation of lifespan brain trajectories, distinguishing longitudinal and cohort effects. Neuroimage, 226, 117596. 10.1016/j.neuroimage.2020.117596

Sripada, C., Rutherford, S., Angstadt, M., Thompson, W. K., Luciana, M., Weigard, A., Hyde, L. H., & Heitzeg, M. (2020). Prediction of neurocognition in youth from resting state fMRI. Mol Psychiatry, 25(12), 3413–3421. 10.1038/s41380-019-0481-6

Stoecklein, S., Hilgendorff, A., Li, M., Forster, K., Flemmer, A. W., Galie, F., Wunderlich, S., Wang, D., Stein, S., Ehrhardt, H., Dietrich, O., Zou, Q., Zhou, S., Ertl-Wagner, B., & Liu, H. (2020). Variable functional connectivity architecture of the preterm human brain: Impact of developmental cortical expansion and maturation. Proc Natl Acad Sci U S A, 117(2), 1201–1206. 10.1073/pnas.1907892117

Sun, L., Zhao, T., Liang, X., Xia, M., Li, Q., Liao, X., Gong, G., Wang, Q., Pang, C., Yu, Q., Bi, Y., Chen, P., Chen, R., Chen, Y., Chen, T., Cheng, J., Cheng, Y., Cui, Z., Dai, Z.,… He, Y. (2025). Human lifespan changes in the brain’s functional connectome. Nat Neurosci, 28(4), 891–901. 10.1038/s41593-025-01907-4

Supekar, K., Musen, M., & Menon, V. (2009). Development of large-scale functional brain networks in children. PLoS biology, 7(7), e1000157.

Sydnor, V. J., Larsen, B., Bassett, D. S., Alexander-Bloch, A., Fair, D. A., Liston, C., Mackey, A. P., Milham, M. P., Pines, A., & Roalf, D. R. (2021). Neurodevelopment of the association cortices: Patterns, mechanisms, and implications for psychopathology. Neuron, 109(18), 2820–2846.

Sydnor, V. J., Larsen, B., Seidlitz, J., Adebimpe, A., Alexander-Bloch, A. F., Bassett, D. S., Bertolero, M. A., Cieslak, M., Covitz, S., Fan, Y., Gur, R. E., Gur, R. C., Mackey, A. P., Moore, T. M., Roalf, D. R., Shinohara, R. T., & Satterthwaite, T. D. (2023). Intrinsic activity development unfolds along a sensorimotor-association cortical axis in youth. Nat Neurosci, 26(4), 638–649. 10.1038/s41593-023-01282-y

Thompson, W. K., Barch, D. M., Bjork, J. M., Gonzalez, R., Nagel, B. J., Nixon, S. J., & Luciana, M. (2019). The structure of cognition in 9 and 10 year-old children and associations with problem behaviors: Findings from the ABCD study’s baseline neurocognitive battery. Dev Cogn Neurosci, 36, 100606. 10.1016/j.dcn.2018.12.004

Tian, Y., & Zalesky, A. (2021). Machine learning prediction of cognition from functional connectivity: Are feature weights reliable? Neuroimage, 245, 118648. 10.1016/j.neuroimage.2021.118648

Vasa, F., Seidlitz, J., Romero-Garcia, R., Whitaker, K. J., Rosenthal, G., Vertes, P. E., Shinn, M., Alexander-Bloch, A., Fonagy, P., Dolan, R. J., Jones, P. B., Goodyer, I. M., consortium, N., Sporns, O., & Bullmore, E. T. (2018). Adolescent Tuning of Association Cortex in Human Structural Brain Networks. Cereb Cortex, 28(1), 281–294. 10.1093/cercor/bhx249

Volkow, N. D., Koob, G. F., Croyle, R. T., Bianchi, D. W., Gordon, J. A., Koroshetz, W. J., Pérez-Stable, E. J., Riley, W. T., Bloch, M. H., & Conway, K. (2018). The conception of the ABCD study: From substance use to a broad NIH collaboration. Developmental cognitive neuroscience, 32, 4–7.

Walhovd, K. B., Fjell, A. M., Giedd, J., Dale, A. M., & Brown, T. T. (2017). Through Thick and Thin: a Need to Reconcile Contradictory Results on Trajectories in Human Cortical Development. Cereb Cortex, 27(2), 1472–1481. 10.1093/cercor/bhv301

Walhovd, K. B., Lovden, M., & Fjell, A. M. (2023). Timing of lifespan influences on brain and cognition. Trends Cogn Sci, 27(10), 901–915. 10.1016/j.tics.2023.07.001

Weintraub, S., Dikmen, S. S., Heaton, R. K., Tulsky, D. S., Zelazo, P. D., Bauer, P. J., Carlozzi, N. E., Slotkin, J., Blitz, D., & Wallner-Allen, K. (2013). Cognition assessment using the NIH Toolbox. Neurology, 80(11_supplement_3), S54-S64.

Xu, Y., Cao, M., Liao, X., Xia, M., Wang, X., Jeon, T., Ouyang, M., Chalak, L., Rollins, N., Huang, H., & He, Y. (2019). Development and Emergence of Individual Variability in the Functional Connectivity Architecture of the Preterm Human Brain. Cereb Cortex, 29(10), 4208–4222. 10.1093/cercor/bhy302

Yan, X., Kong, R., Xue, A., Yang, Q., Orban, C., An, L., Holmes, A. J., Qian, X., Chen, J., & Zuo, X.-N. (2023). Homotopic local-global parcellation of the human cerebral cortex from resting-state functional connectivity. Neuroimage, 273, 120010.

Yeo, B. T., Krienen, F. M., Sepulcre, J., Sabuncu, M. R., Lashkari, D., Hollinshead, M., Roffman, J. L., Smoller, J. W., Zollei, L., Polimeni, J. R., Fischl, B., Liu, H., & Buckner, R. L. (2011). The organization of the human cerebral cortex estimated by intrinsic functional connectivity. J Neurophysiol, 106(3), 1125–1165. 10.1152/jn.00338.2011

Zhang, S., Larsen, B., Sydnor, V. J., Zeng, T., An, L., Yan, X., Kong, R., Kong, X., Gur, R. C., Gur, R. E., Moore, T. M., Wolf, D. H., Holmes, A. J., Xie, Y., Zhou, J. H., Fortier, M. V., Tan, A. P., Gluckman, P., Chong, Y. S.,… Yeo, B. T. T. (2024). In vivo whole-cortex marker of excitation-inhibition ratio indexes cortical maturation and cognitive ability in youth. Proc Natl Acad Sci U S A, 121(23), e2318641121. 10.1073/pnas.2318641121

Zhi, D., Jiang, R., Pearlson, G., Fu, Z., Qi, S., Yan, W., Feng, A., Xu, M., Calhoun, V., & Sui, J. (2024). Triple Interactions Between the Environment, Brain, and Behavior in Children: An ABCD Study. Biol Psychiatry, 95(9), 828–838. 10.1016/j.biopsych.2023.12.019

