## supplementary_information for "Convergent and Divergent Brain–Cognition Development"

Contents

**Supplementary Methods**

Mathematical relationship between the reliability of FC change and reliability of cross-sectional FC

**Supplementary Figures**

Figure S1 Illustration of how cross-sectional and longitudinal analyses can lead to divergent estimates

Figure S2 Enhanced FC-cognition relationship during development

Figure S3 Models trained on baseline FC to predict baseline cognition improve in accuracy when applied to Year 2 FC and Year 2 cognition

Figure S4 Models trained on baseline FC to predict baseline cognition improve in accuracy when applied to Year 2 FC and Year 2 cognition, even after controlling for head motion.

Figure S5 Baseline FC is more predictive of cognition at Year 2 than longitudinal FC change, even accounting for reliability differences

Figure S6 Baseline FC is more predictive of cognition at Year 2 than rate of FC change

Figure S7 Baseline FC and rate of FC change weakly predict rate of cognitive change

Figure S8 Convergent and divergent predictive network features (PNFs) between cross-sectional and longitudinal estimates of FC-cognition relationship for the eight cognitive measures

**Supplementary References**

**Mathematical relationship between the reliability of FC change and the reliability of cross-sectional FC**

Here we provide a detailed derivation linking the reliability of FC change to the reliability of FC at two timepoints (Rogosa & Willett, 1983).

Denote an FC edge at time point 1 as $FC_1$ and that same FC edge at time point 2 as $FC_2$. We can decompose $FC_1$ and $FC_2$ into 2 components:

$$\begin{aligned} \mathrm{FC}_1 &= T_1 + \epsilon_1 \\ \mathrm{FC}_2 &= T_2 + \epsilon_2 \end{aligned}$$

where $T_1$ and $T_2$ are the true noise-free components of the FCs and $\epsilon$ are noises.

Define the reliability of $FC_1$ as $R_1 = \frac{V(T_1)}{V_1}$. Similarly, $R_2 = \frac{V(T_2)}{V_2}$.

In general, for a given random variable $X$, we have $V_X = V(T_X) + V(\epsilon_X)$. Therefore,

$$\begin{aligned} V(\epsilon_X) &= V_X - V(T_X) \\ &= V_X - R_X V_X \\ &= (1 - R_X)V_X \end{aligned}$$

More specifically, $V(\epsilon_1) = (1 - R_1)V_1$ and $V(\epsilon_2) = (1 - R_2)V_2$.

Our question is to estimate the reliability of $\mathrm{FC}_D = \mathrm{FC}_1 - \mathrm{FC}_2$ or $R_D$.

First,

$$\mathrm{FC}_D = \mathrm{FC}_1 - \mathrm{FC}_2 = T_1 - T_2 + \epsilon_1 - \epsilon_2$$

Then,

$$\begin{aligned} R_D &= \frac{V(T_D)}{V_D} \\ &= \frac{V(T_1) + V(T_2) - 2\mathrm{Cov}(T_1, T_2)}{V(T_1) + V(T_2) - 2\mathrm{Cov}(T_1, T_2) + V(\epsilon_1) + V(\epsilon_2) - \cancel{2\mathrm{Cov}(\epsilon_1, \epsilon_2)}} \\ &= \frac{V(T_1) + V(T_2) - 2\rho_T\sqrt{V(T_1)V(T_2)}}{V(T_1) + V(T_2) - 2\rho_T\sqrt{V(T_1)V(T_2)} + V(\varepsilon_1) + V(\varepsilon_2)} \\ &= \frac{R_1V_1 + R_2V_2 - 2\rho_T\sqrt{R_1V_1R_2V_2}}{R_1V_1 + R_2V_2 - 2\rho_T\sqrt{R_1V_1R_2V_2} + (1 - R_1)V_1 + (1 - R_2)V_2} \\ &= \frac{R_1V_1 + R_2V_2 - 2\rho_T\sqrt{R_1V_1R_2V_2}}{V_1 + V_2 - 2\rho_T\sqrt{R_1V_1R_2V_2}} \qquad (1) \end{aligned}$$

where $\rho_T$ is the correlation between $T_1$ and $T_2$. Define ρ as the correlation between $FC_1$ and $FC_2$. Since $\rho = \rho_T\sqrt{R_1R_2}$, equation (1) becomes

$$R_D = \frac{R_1V_1 + R_2V_2 - 2\rho\sqrt{V_1V_2}}{V_1 + V_2 - 2\rho\sqrt{V_1V_2}} \qquad (2)$$

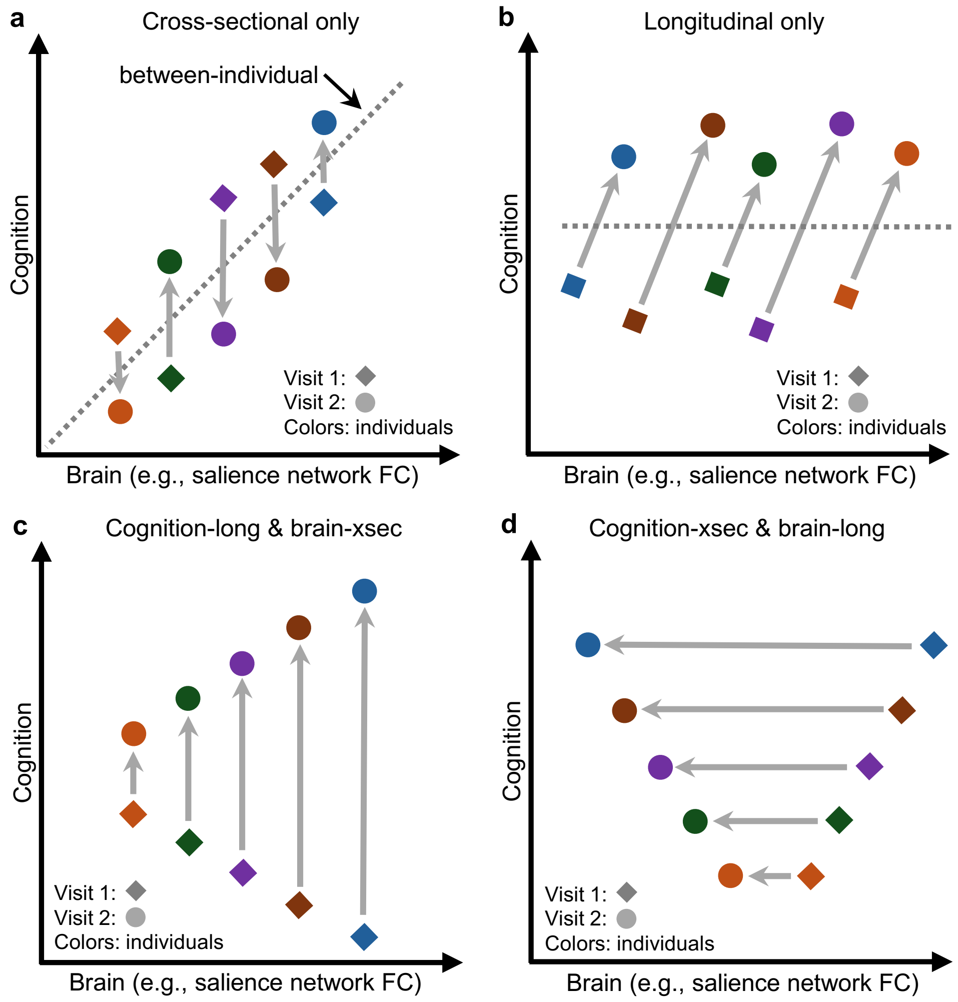


**Figure S1.** Illustration of how cross-sectional and longitudinal analyses can lead to divergent estimates. (a) A cross-sectional brain–cognition relationship exists, but not longitudinal one. Individual-averaged brain measures are associated with individual-averaged cognitive measures, but within-individual values are not associated. (b) A longitudinal brain–cognition relationship exists, but not cross-sectional one. Within-individual change in brain measures and cognitive measures are associated, but individual-averaged values are not associated. (c) Cross-sectional brain measures are associated with longitudinal cognitive change. Individual-averaged brain measures are associated with within-individual changes in cognition, but within-individual brain measures are not informative. (d) Longitudinal brain measures are associated with individual-averaged cognitive measures. Within-individual changes in brain measures are associated with individual-averaged cognition measures, but individual-averaged brain measures are not informative.

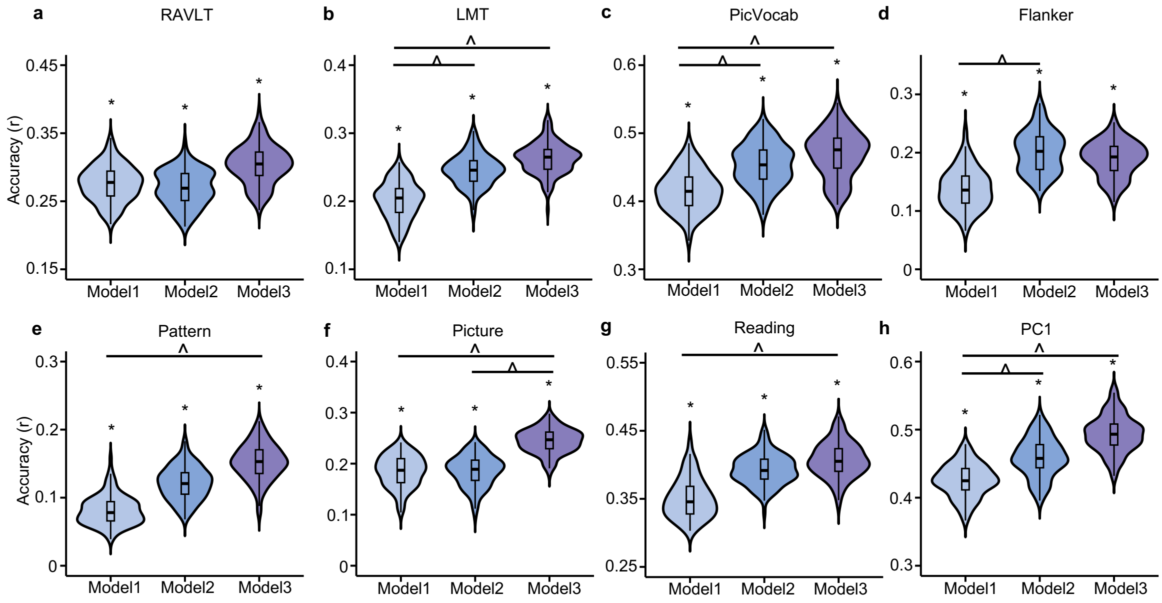


**Figure S2**. Enhanced FC-cognition relationship during development. (A-H) Comparison of prediction accuracies across the three models for the eight cognitive measures. Model 1 predicted baseline cognition using baseline FC (FCY0 → CogY0). Model 2 predicted Year 2 cognition using baseline FC (FCY0 → CogY2). Model 3 predicted Year 2 cognition using Year 2 FC (FCY2 → CogY2). Each value in the violin plot represents the accuracy (r) for a single cross-validation fold. Asterisks (*) denote above chance prediction after multiple comparisons correction (FDR $q < 0.05$). Carets (^) denote statistically significant differences between models based on the corrected resampled t-test (FDR $q < 0.05$). Note: Y-axes differ across panels to enhance visibility and emphasize the performance of models within each task. Comparisons across tasks should be made with caution due to varying scales. RAVLT: Rey Auditory Verbal Learning Test (verbal memory); LMT: Little Man Task (spatial reasoning); PicVocab: Picture Vocabulary Task (vocabulary); Flanker: Flanker Task (executive function); Pattern: Pattern Comparison Processing Speed Test (processing speed); Picture: Picture Sequence Memory Test (episodic memory); Reading: Oral Reading Recognition Task (reading ability). PC1: the first principal component of the above seven cognitive measures.

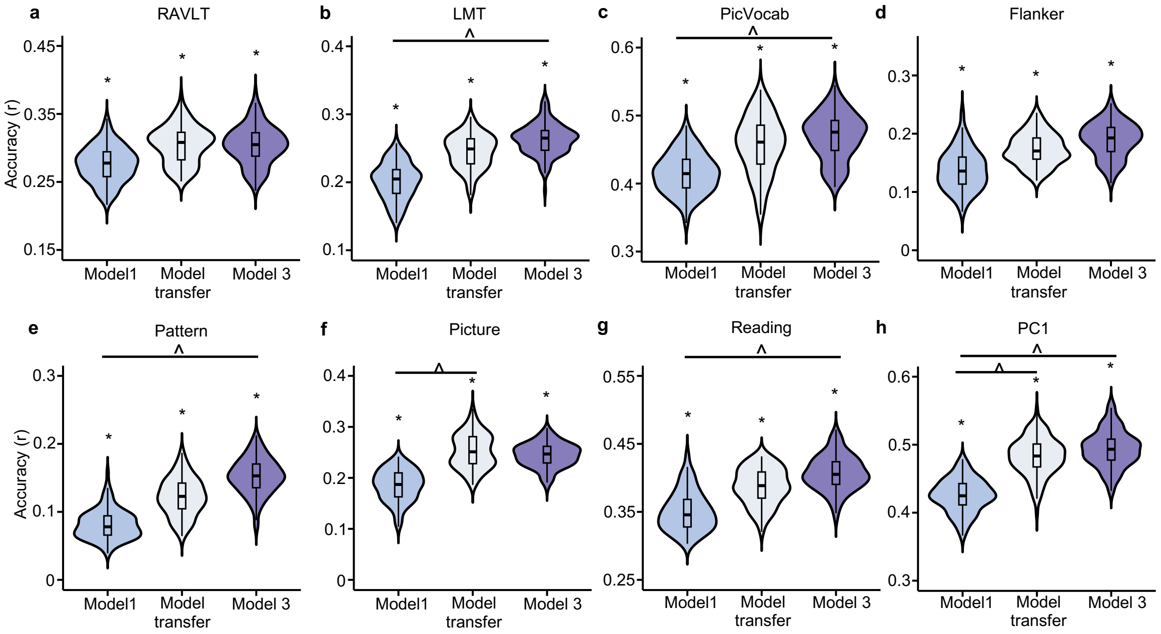


**Figure S3.** Models trained on baseline FC to predict baseline cognition improve in accuracy when applied to Year 2 FC and Year 2 cognition. Each plot corresponds to a different cognitive measure. "Model 1" are the models obtained by training baseline FC to predict baseline cognition (FCY0 → CogY0). These models were then used to predict Y2 cognition from Year 2 FC, which we refer to as "Model transfer". Finally, for reference, "Model 3" are the models obtained by training Y2 FC to predict Y2 cognition (FCY2 → CogY2). Asterisks (*) denote above chance prediction after multiple comparisons correction (FDR $q < 0.05$). Carets (^) denote statistically significant differences between models based on the corrected resampled t-test (FDR $q < 0.05$). Note: Y-axes differ across panels to enhance visibility and emphasize the performance of models within each task. Comparisons across tasks should be made with caution due to varying scales. RAVLT: Rey Auditory Verbal Learning Test (verbal memory); LMT: Little Man Task (spatial reasoning); PicVocab: Picture Vocabulary Task (vocabulary); Flanker: Flanker Task (executive function); Pattern: Pattern Comparison Processing Speed Test (processing speed); Picture: Picture Sequence Memory Test (episodic memory); Reading: Oral Reading Recognition Task (reading ability). PC1: the first principal component of the above seven cognitive measures.

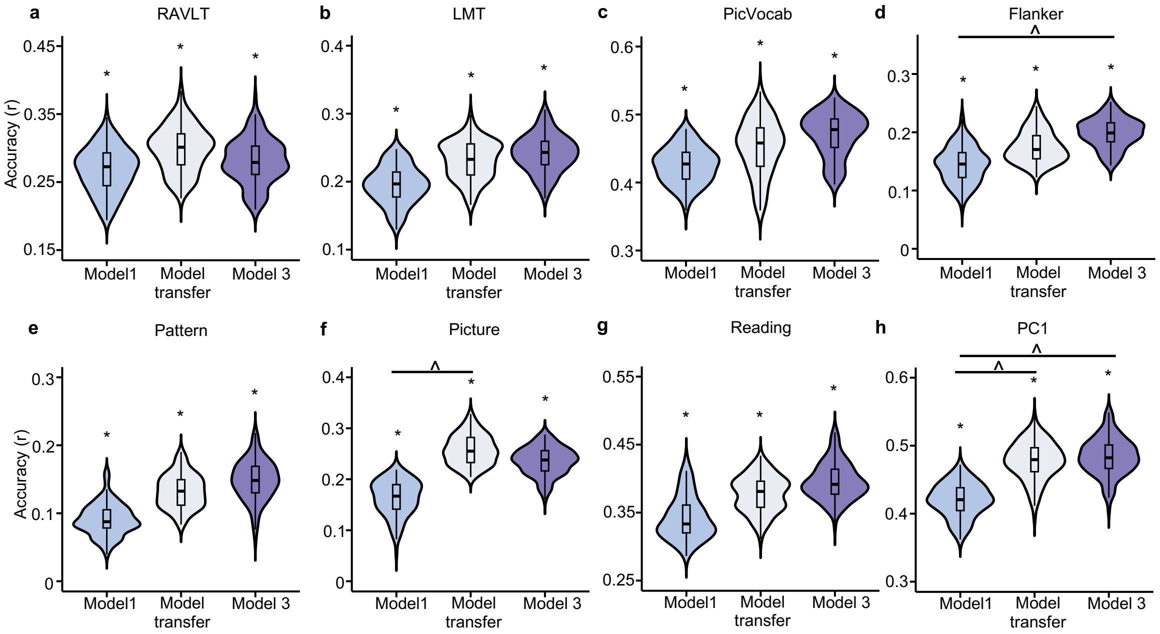


**Figure S4.** Models trained on baseline FC to predict baseline cognition improve in accuracy when applied to Year 2 FC and Year 2 cognition, even after controlling for head motion. Head motion was controlled by selecting a subset of participants (n = 2642) that with no significant difference in mean frame-wise displacement across the two timepoints (p = 0.18). Each plot corresponds to a different cognitive measure. "Model 1" are the models obtained by training baseline FC to predict baseline cognition (FCY0 → CogY0). These models were then used to predict Year 2 cognition from Year 2 FC, which we refer to as "Model transfer". Finally, for reference, "Model 3" are the models obtained by training Y2 FC to predict Year 2 cognition (FCY2 → CogY2). Asterisks (*) denote above chance prediction after multiple comparisons correction (FDR q < 0.05). Carets (^) denote statistically significant differences between models based on the corrected resampled t-test (FDR q < 0.05). Note: Y-axes differ across panels to enhance visibility and emphasize the performance of models within each task. Comparisons across tasks should be made with caution due to varying scales. RAVLT: Rey Auditory Verbal Learning Test (verbal memory); LMT: Little Man Task (spatial reasoning); PicVocab: Picture Vocabulary Task (vocabulary); Flanker: Flanker Task (executive function); Pattern: Pattern Comparison Processing Speed Test (processing speed); Picture: Picture Sequence Memory Test (episodic memory); Reading: Oral Reading Recognition Task (reading ability). PC1: the first principal component of the above seven cognitive measures.

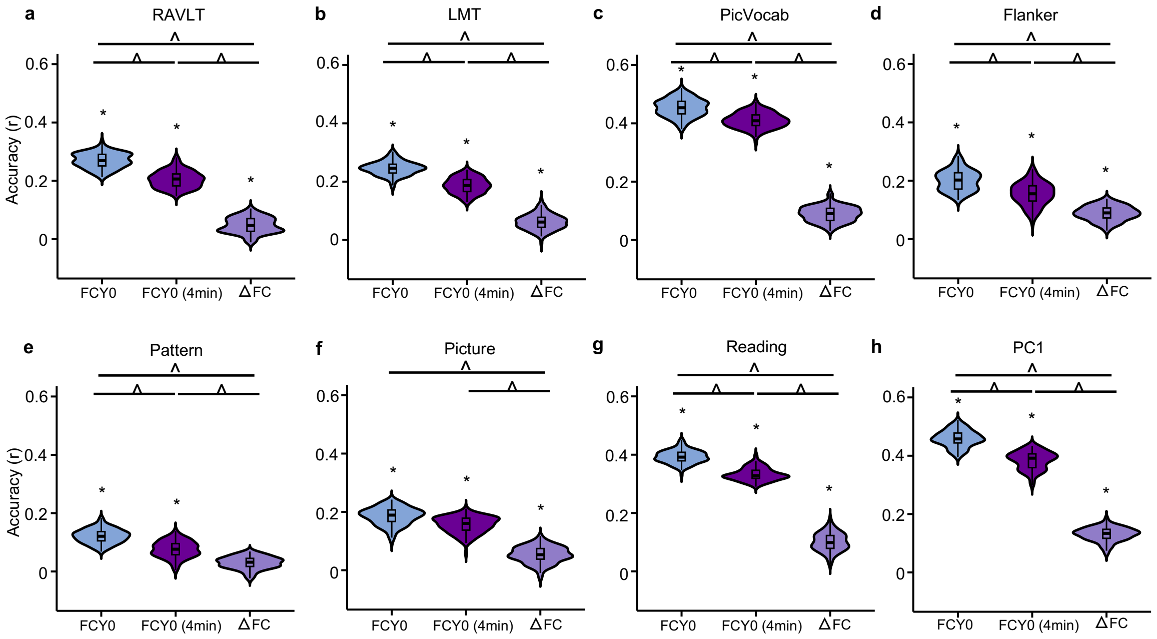


**Figure S5.** Baseline FC is more predictive of cognition at Year 2 than longitudinal FC change, even accounting for reliability differences. Each panel corresponds to the prediction accuracy of a different cognitive measure at Year 2. "FCY0" are the models obtained by training baseline FC to predict Year 2 cognition (same as model 2 in Figure 4). "FC (4min)" uses baseline FC computed from the first 4 minutes of fMRI data to predict Year 2 cognition. "ΔFC" are the models obtained by using FC change (between Year 2 and baseline) to predict cognition at Year 2. Asterisks (*) denote above chance prediction after multiple comparisons correction (FDR $q < 0.05$). Carets (^) denote statistically significant differences between models based on the corrected resampled t-test (FDR $q < 0.05$). RAVLT: Rey Auditory Verbal Learning Test (verbal memory); LMT: Little Man Task (spatial reasoning); PicVocab: Picture Vocabulary Task (vocabulary); Flanker: Flanker Task (executive function); Pattern: Pattern Comparison Processing Speed Test (processing speed); Picture: Picture Sequence Memory Test (episodic memory); Reading: Oral Reading Recognition Task (reading ability). PC1: the first principal component of the above seven cognitive measures.

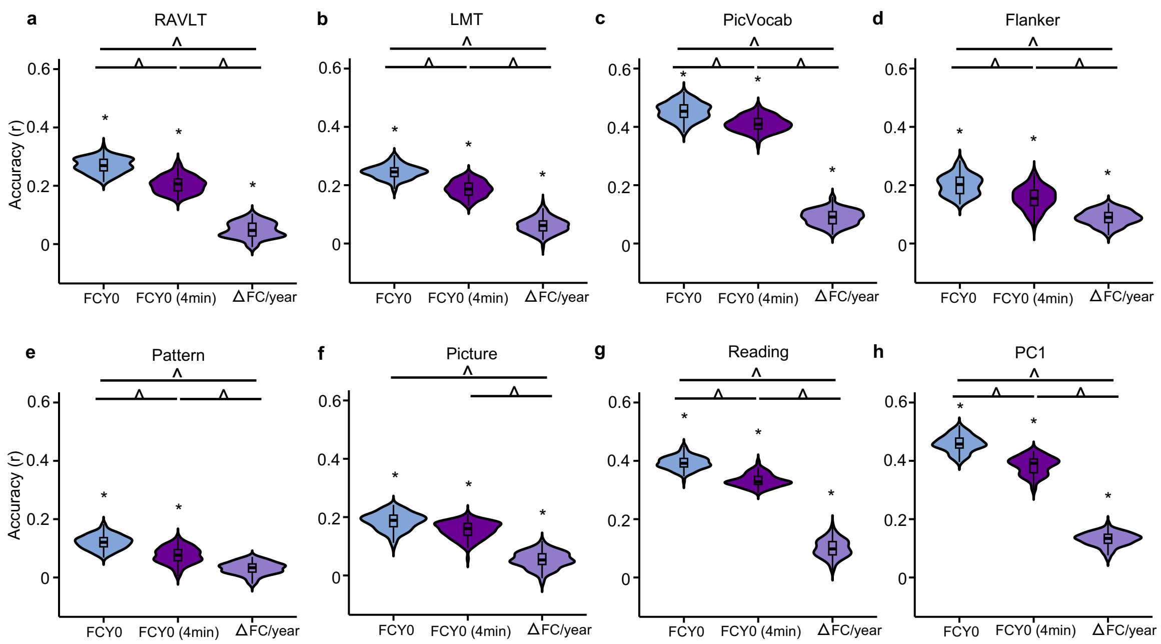


**Figure S6.** Baseline FC is more predictive of cognition at Year 2 than rate of FC change. Rate of FC change (ΔFC/year) was used instead of the FC change (delta) in this validation. Each panel corresponds to the prediction accuracy of a different cognitive measure at Year 2. "FCY0" are the models obtained by training baseline FC to predict Year 2 cognition (same as model 2 in Figure 4). "FC (4min)" uses baseline FC computed from the first 4 minutes of fMRI data to predict Year 2 cognition. "ΔFC/year" are the models obtained by using rate of FC change (between Year 2 and baseline) to predict cognition at Year 2. Asterisks (*) denote above chance prediction after multiple comparisons correction (FDR $q < 0.05$). Carets (^) denote statistically significant differences between models based on the corrected resampled t-test (FDR $q < 0.05$). RAVLT: Rey Auditory Verbal Learning Test (verbal memory); LMT: Little Man Task (spatial reasoning); PicVocab: Picture Vocabulary Task (vocabulary); Flanker: Flanker Task (executive function); Pattern: Pattern Comparison Processing Speed Test (processing speed); Picture: Picture Sequence Memory Test (episodic memory); Reading: Oral Reading Recognition Task (reading ability). PC1: the first principal component of the above seven cognitive measures.

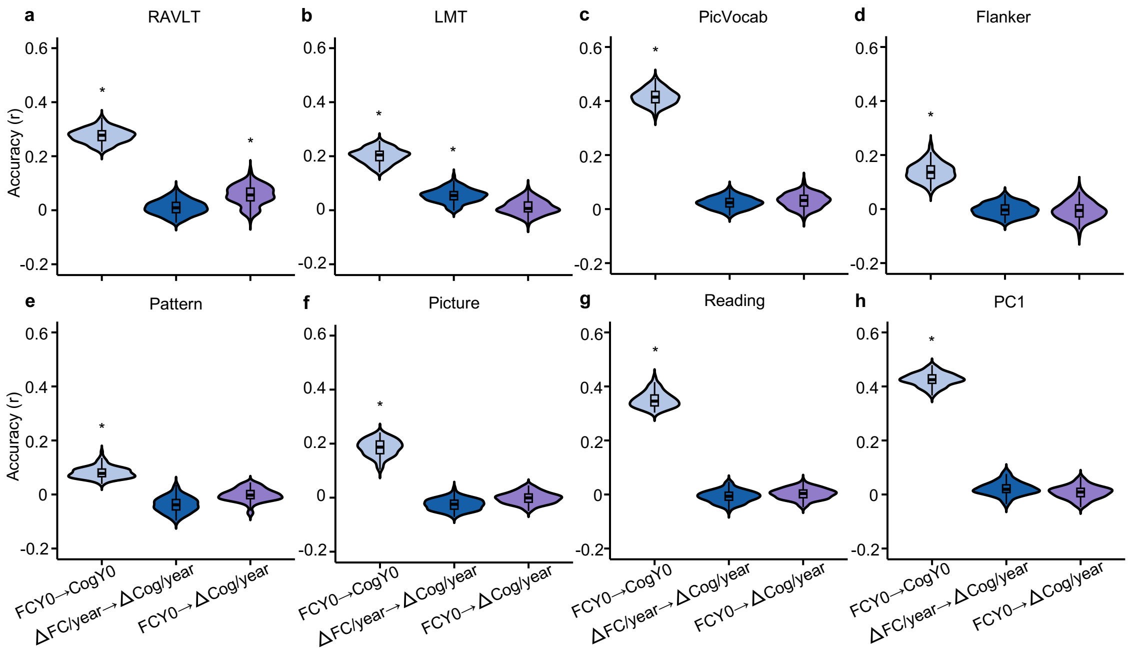


**Figure S7.** Baseline FC and rate of FC change weakly predict rate of cognitive change. Rate of FC change (ΔFC/year) and rate of cognitive change (ΔCog/year) were used instead of the FC and cognitive change (delta) in this validation. Each panel corresponds to the prediction accuracy of a different cognitive measure. Within each panel, baseline FC and rate of FC change (ΔFC/year) were used to predict rate of cognitive change (ΔCog/year). Predictions of baseline cognition from baseline FC (FCY0 → CogY0) were also shown for reference. Asterisks (*) denote above chance prediction after multiple comparisons correction (FDR $q < 0.05$). RAVLT: Rey Auditory Verbal Learning Test (verbal memory); LMT: Little Man Task (spatial reasoning); PicVocab: Picture Vocabulary Task (vocabulary); Flanker: Flanker Task (executive function); Pattern: Pattern Comparison Processing Speed Test (processing speed); Picture: Picture Sequence Memory Test (episodic memory); Reading: Oral Reading Recognition Task (reading ability). PC1: the first principal component of the above seven cognitive measures.

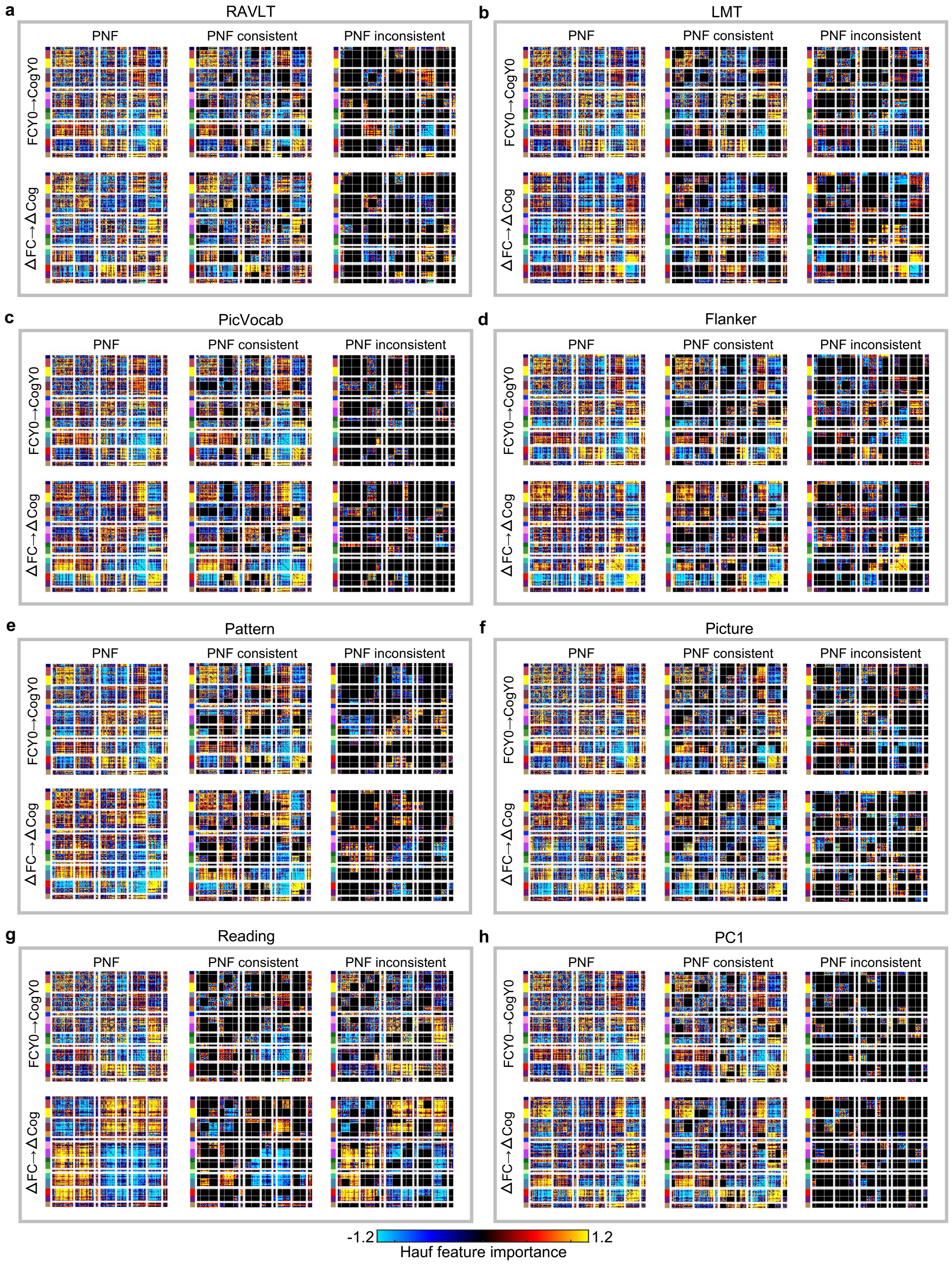


**Figure S8.** Convergent and divergent predictive network features (PNFs) between cross-sectional and longitudinal estimates of FC-cognition relationships for the eight cognitive measures. Each grey box represents one cognitive measure. Within each box, the first row corresponds to the cross-sectional model (FCY0 → CogY0), and the second row corresponds to the longitudinal model (ΔFC → ΔCog). The PNF column displays the predictive network features for each model. The PNF consistent column highlights network blocks where the average PNFs have the same sign in both models, indicating convergence. The PNF inconsistent column highlights blocks where the average PNFs have opposite signs, indicating divergence. RAVLT: Rey Auditory Verbal Learning Test (verbal memory); LMT: Little Man Task (spatial reasoning); PicVocab: Picture Vocabulary Task (vocabulary); Flanker: Flanker Task (executive function); Pattern: Pattern Comparison Processing Speed Test (processing speed); Picture: Picture Sequence Memory Test (episodic memory); Reading: Oral Reading Recognition Task (reading ability). PC1: the first principal component of the above seven cognitive measures. For visualization purposes, each predictive network feature matrix was normalized by dividing all values by the standard deviation of the entire matrix.

### References


Rogosa, D. R., & Willett, J. B. (1983). Demonstrating the Reliability of the Difference Score in the Measurement of Change. *Journal of Educational Measurement*, *20*, 335-343. https://www.jstor.org/stable/1434950
